# Transcriptional Diversity of Medium Spiny Neurons in the Primate Striatum

**DOI:** 10.1101/2020.10.25.354159

**Authors:** Jing He, Michael Kleyman, Jianjiao Chen, Aydin Alikaya, Kathryn M. Rothenhoefer, Bilge Esin Ozturk, Morgan Wirthlin, Kenneth Fish, Leah C. Byrne, Andreas R. Pfenning, William R. Stauffer

## Abstract

The striatum is the interface between dopamine reward signals and cortico-basal ganglia circuits that mediate diverse behavioral functions. Medium spiny neurons (MSNs) constitute the vast majority of striatal neurons and are traditionally classified as direct- or indirect-pathway neurons. However, that traditional model does not explain the anatomical and functional diversity of MSNs. Here, we defined molecularly distinct MSN types in the primate striatum, including (1) dorsal striatum MSN types associated with striosome and matrix compartments, (2) ventral striatum types associated with the nucleus accumbens shell and olfactory tubercle, and (3) an MSN-like type restricted to μ-opioid receptor rich islands in the ventral striatum. These results lay the foundation for achieving cell type-specific transgenesis in the primate striatum and provide a blueprint for investigating circuit-specific processing.

## INTRODUCTION

The striatum serves as the major input nucleus for the basal ganglia and the principal neural interface between dopamine reward signals and cortico-basal ganglia-thalamo-cortical circuits. Medium Spiny Neurons (MSNs), also known as Spiny Projection Neurons (SPNs), account for the vast majority of all striatal neurons (DiFiglia et al., 1976). D1-MSNs express dopamine receptor type 1 (*DRD1*) and their monosynaptic projections to the basal ganglia output nuclei form the “direct pathway” (Parent and Hazrati, 1995). D2-MSNs express dopamine receptor type 2 (*DRD2*) and form the “indirect pathway” via di-synaptic projections to the basal ganglia output nuclei (Parent and Hazrati, 1995). Activity in the direct and indirect pathways produces, broadly, opposing effects on thalamo-cortical projections. This cell type-specific circuit model has been crucial to understanding the role of the basal ganglia in the control of movement and the mechanisms of Parkinson’s disease (Albin et al., 1989; DeLong and Wichmann, 2015). However, this model does not account for the anatomical and functional variations between the distinct anatomical compartments in the striatum, nor does it account for the striatum’s diverse behavioral roles.

In the dorsal striatum (DS), direct and indirect pathway information processing occurs within largely segregated neurochemical compartments known as striosome (patch) and matrix (Graybiel and Ragsdale, 1978). Functionally, these compartments are thought to participate in limbic and sensorimotor functions, respectively (Gerfen, 1984; Graybiel and Ragsdale, 1978; Hong et al., 2019). The ventral striatum (VS), which includes the nucleus accumbens (NAc), olfactory tubercle (OT), and interface islands, is crucial for rewards, learning and emotional responses (Daunais et al., 2001; Haber and McFarland, 1999; Heimer and Wilson, 1975; Voorn et al., 1996). Moreover, the NAc itself is divided into core and shell territories that contribute differentially to reward processing, action selection, and decision making (Floresco, 2015). Single cell technologies that classify cell types according to their overall gene expression profiles provide powerful and quantitative methods for investigating cell type heterogeneity (Tang et al., 2009). Single cell and single nucleus RNA sequencing (scRNA-Seq and snRNA-Seq, respectively) has revealed new types of MSNs and striatal interneurons (Gokce et al., 2016; Krienen et al., 2020; Martin et al., 2019; Munoz-Manchado et al., 2018; Saunders et al., 2018; Stanley et al., 2020). Moreover, these technologies are providing novel insights into cell type-specific mechanisms for diseases involving the striatum, including drug addiction (Savell et al., 2020) and Huntington’s disease (Lee et al., 2020). Despite these advances, we know neither the extent of MSN diversity in the primate striatum, nor how that diversity corresponds to the anatomical or neurochemical divisions of the highly articulated primate brain.

The close phylogenetic relationship and the high degree of homology between human and non-human primate (NHP) brains, genes, and behaviors make it evident that NHP studies are indispensable for understanding the neuronal substrates of human behavior as well as neurological, neurodegenerative, and psychiatric diseases (Izpisua Belmonte et al., 2015). Here, we set out to identify molecularly distinct types of MSNs in Rhesus macaque monkeys. We used snRNA-Seq and Fluorescent In-Situ Hybridization (FISH) to characterize the transcriptional and anatomical diversity of MSNs and closely related neurons. Our results indicate that MSN types, each of which was distinguishable by a specific set of differentially expressed genes, reflect the major anatomical and neurochemical divisions, including striosome and matrix compartments of the dorsal striatum, the core and shell regions of the NAc, and the ventral interface islands including the Islands of Calleja (ICj) and Neurochemically Unique Domains in the Accumbens and Putamen (NUDAPs) (Voorn et al., 1996). The cell type-specific gene expression patterns provide insights into MSN functions and indicate potential molecular access points for cell type-specific applications of genetically coded tools to primate brains in scientific or translational settings.

## RESULTS

### Major Cell Classes in the Primate Striatum

To investigate the cell type-specific transcriptional architecture of the NHP striatum, we micro-dissected the caudate (Cd), putamen (Pt), and ventral striatum (VS) from coronal sections (Figure 1A) of two monkeys.Using these samples, we prepared single nuclei suspensions, performed droplet-based RNA capture, and sequenced the resulting barcoded cDNA libraries (STAR Methods). To optimally align and count the transcripts, we re-annotated the Rhesus macaque reference genome (rheMac10) using a “liftOver” of the human transcriptome annotation; a process that improves comparisons across species (Zhu et al., 2014). The custom annotation increased the reads confidently mapped to the genome by more than 10% (p = 1.9 × 10^−10^, paired t-test, Figure S1A). We used Seurat to integrate the datasets from both monkeys, and side by side comparison of the nuclei from both subjects confirmed successful integration (Figure 1B). After removing technical artifacts, we recovered the gene expression profiles from approximately 80,000 nuclei in two monkeys (F and P). These profiles were used for all subsequent analysis.

**Figure 1.**
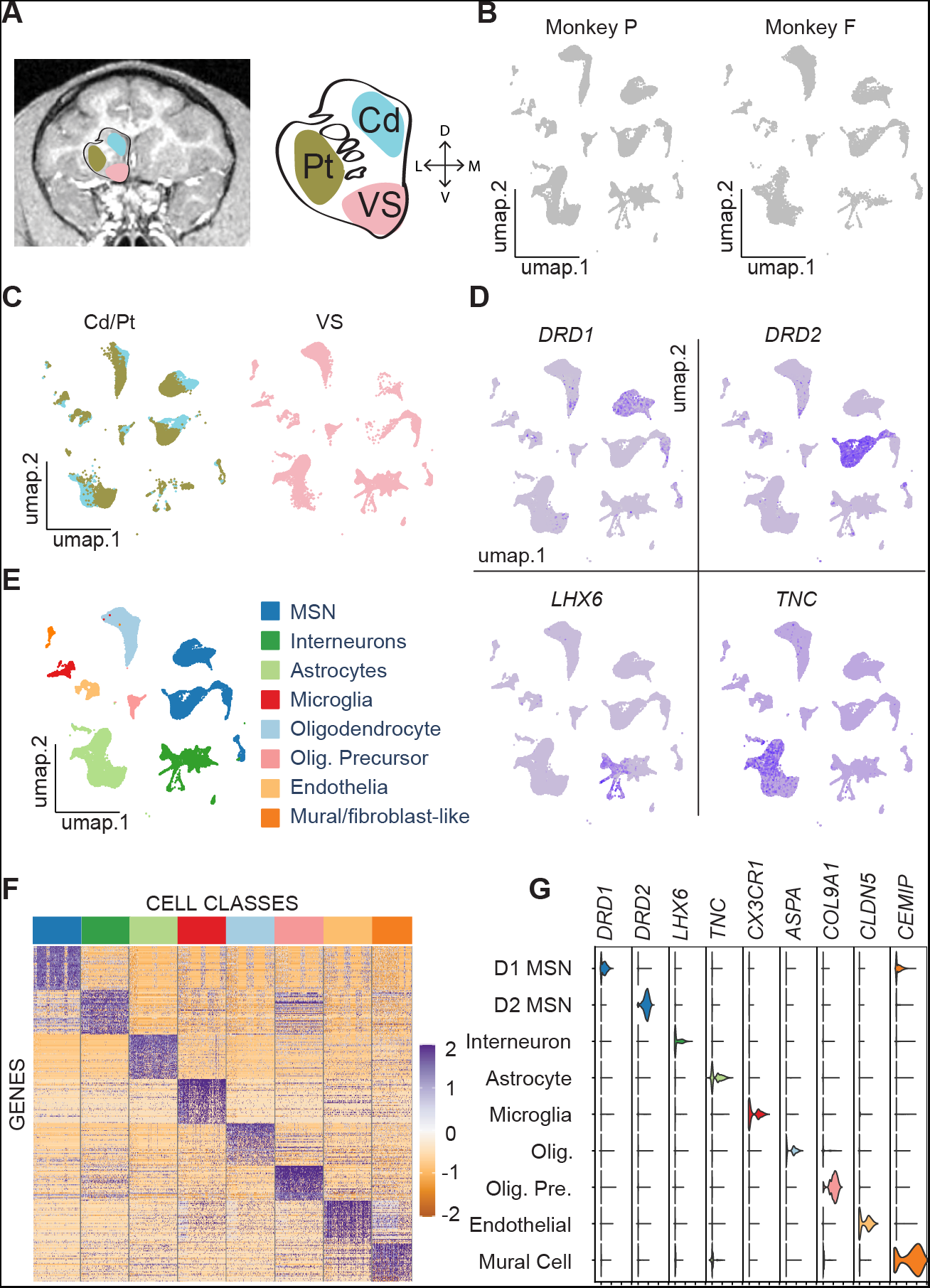
Cell Type Taxonomy in Primate Striatum. (A)MRI image of a Rhesus macaque coronal brain section (left) showing three striatal regions labeled by three patches (cyan, caudate; brown, putamen; pink, ventral striatum). Schematic striatum (middle) marked by Cd (caudate), Pt (putamen), and VS (ventral striatum). The right axis shows dorsal (D), ventral (V), lateral (L), and medial (M) directions. (B)UMAP visualizations of the samples from the two subjects. (C)UMAP visualizations of striatal nuclei colored by the three regions. The color scheme for these regions is the same as in A. (D)Feature plots of canonical neuronal and astrocyte marker gene expression in striatal nuclei. (E)UMAP visualization colored according to eight major classes in the NHP. (F)Heat map of differentially expressed genes. Color bar at the top corresponds to the major classes identified in E. (G)Violin plots of distributions of marker gene expression across nine clusters, with MSNs being divided into D1- and D2-MSNs. See also Figure S1.

To identify the major cell classes in the striatum, we clustered the nuclei based on their gene expression counts (see STAR Methods). Each major cluster had nuclei derived from the Cd, Pt, and VS, indicating broad similarities between the cell type compositions of each region (Figure 1C). We used feature plots that showed the expression of well-known marker genes including *DRD1*, *DRD2*, *LHX6*, to identify broad neuronal classes including D1-MSNs, D2-MSN, and interneurons, respectively (Figure 1D). Similarly, we identified astrocytes, oligodendrocytes, and other non-neuronal cell types (Figures 1D and S1B). Each major cell class was signified by groups of differentially expressed genes that were specifically enriched in that cluster (Figures 1E, 1F and Table S1). We compared the intersection of differentially expressed genes from each cluster with orthologous marker gene lists from several databases (Gokce et al., 2016; Saunders et al., 2018; Zhang et al., 2014). This comparison confirmed the identity of the clusters and their correspondence to the major cell classes in the striatum (p < 0.05, Benjamini-Hochberg corrected, Hypergeometric tests). Violin plots showing the expression levels of marker genes in each major class confirmed the basic validity of our experimental and analytic methods (Figure 1G). Together, these results provide a broad transcriptional catalogue of cell classes in the NHP striatum and a rich dataset for future exploration.

### Transcriptional Diversity of Medium Spiny Neurons

According to our primary interest, we sought to isolate and analyze MSN nuclei. We identified all neurons according to the expression of *RBFOX3*, the gene that codes for the neuron-specific marker NeuN (Figure S1C). Within this population that included interneurons, MSNs were isolated according to the expression of several well-known marker genes including *PP1R1B*, *BCL11B*, and *PDE1B* (Figures S1D-S1F) (Arlotta et al., 2008; Xie et al., 2006). Differential gene analysis of the isolated MSNs versus all other cell types revealed several other general MSN markers, including *KIAA1211L*, *PDE2A, SLIT3*, and *NGEF* (Figures S1G-S1J).

We clustered MSNs at a resolution that putatively reflected the major functional divisions in the striatum, including dorsal and ventral striatum (DS and VS, respectively), striosome and matrix compartments, and core and shell territories (STAR Methods). This resulted in nine clusters (Figure 2A). The proportion of nuclei from monkey P and F in each cluster (Figure S1N), and the number of genes expressed in each cluster was roughly similar (Figure S1O). The feature plots of canonical D1- and D2-MSN markers showed that the clusters mostly maintained clear separation between *DRD1*- and *DRD2*-expressing nuclei (Figure 2B). To identify and annotate the clusters, we used previously described marker genes for striatal neurons (Crittenden and Graybiel, 2011; Martin et al., 2019; Smith et al., 2016) and subsequent FISH labeling of markers identified from differential gene analysis (Figures 2C, and 3-7). The clusters corresponded to D1- and D2-MSNs in the matrix, D1- and D2-MSNs in striosomes, D1- and D2-MSNs in the NAc shell and olfactory tubercle (OT), and MSN-like neurons located in the interface islands (Figure 2D). One small cluster was a D1/D2-hybrid and likely corresponded to a novel MSN type recently described in rodents (D1H or eccentric-SPN) (Figures 2D and S2A-S2C) (Saunders et al., 2018; Stanley et al., 2020).

**Figure 2.**
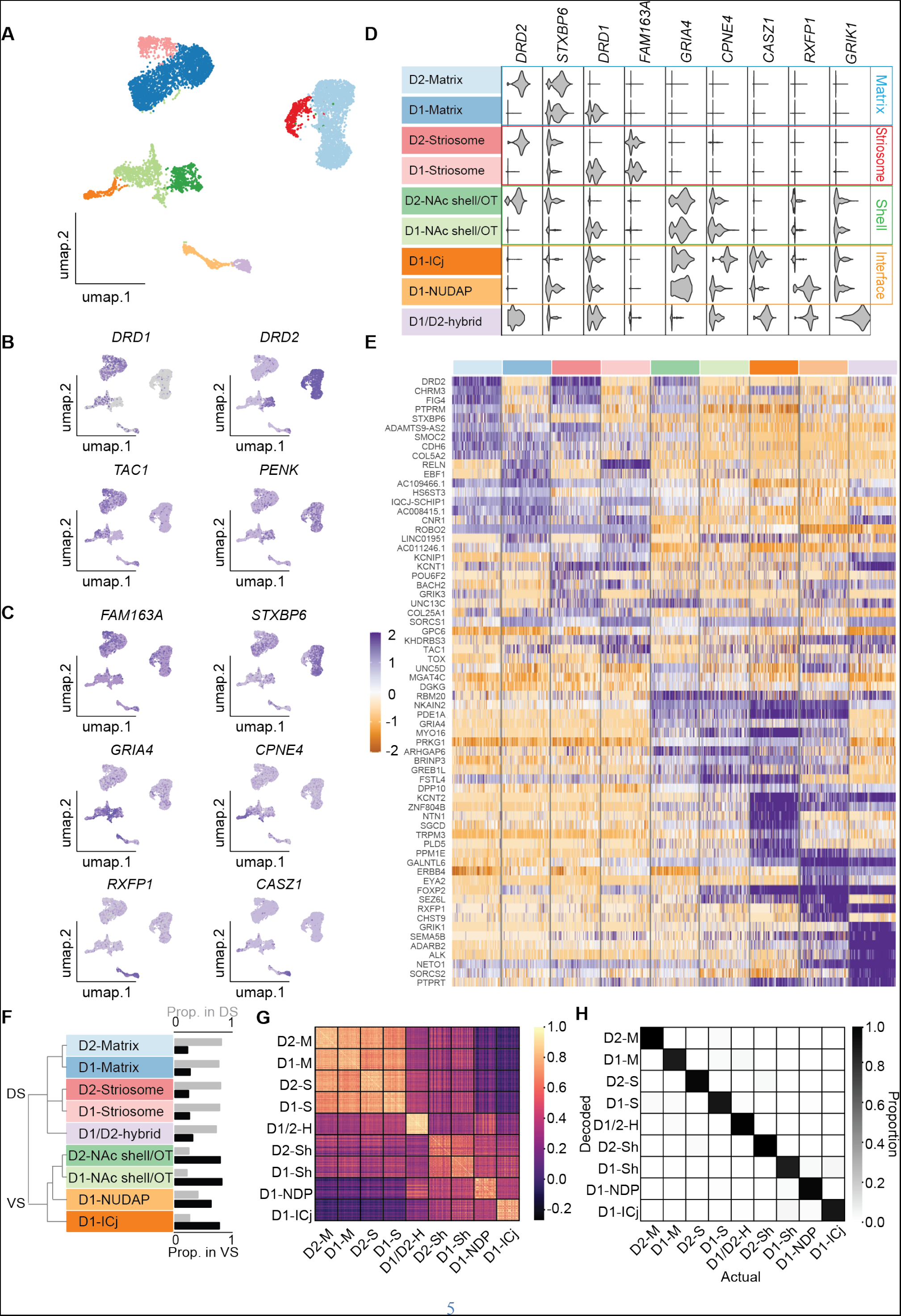
MSN Transcriptional Diversity. (A)UMAP projection of MSN nuclei. Each dot represents a nucleus and the colors represent the different MSN types. (B)Feature plots for the expression of *DRD1*, *TAC1*, *DRD2*, *PENK* show the separation of D1- and D2-MSNs. (C)Feature plots showing marker genes most enriched in each cluster. (D)(left) MSN type identifications colored according to A. (right) Violin plots showing cell type and compartment specific marker gene expression. (E)Heatmap showing the top ten most enriched genes in each MSN type. Colored bar at the top corresponds to the colors in A. (F)(left) Hierarchical clustering of the nine clusters arranged cell type according to anatomical distribution. (right) Proportion of cells in the DS and VS for each cluster. (G)Heat map showing the cosine similarity between cells within and between the nine types of MSNs. (H)The accuracy rate between SCCAF decoded cell type and actual cell type. See also Figure S1 and S2.

Each MSN cluster was signified by groups of differentially expressed genes that were specifically enriched in that cluster (Figure 2E and Table S2). Complete linkage hierarchical clustering arranged the nine cell types such that the first split occurred between DS and VS clusters and reflected roughly the proportions of nuclei derived from the DS and VS within each cluster (Figure 2F). Next, the hierarchical clustering algorithm split the clusters according to major anatomical and functional divisions, including striosome and matrix, and finally into division specific D1-and D2-MSN clusters (Figure 2F). We validated this clustering using cosine similarity between individual nuclei (STAR Methods). For each of the nine clusters, nuclei within a cluster were far more similar to each other, than to nuclei in other clusters (p < 0.0001, Permutation test, STAR Methods) (Figure 2G). Moreover, nuclei within clusters derived mostly from the DS were more similar to each other than VS derived nuclei, and the reverse was also true (p < 0.0001, Permutation tests, STAR Methods). These results suggest that the major functional and anatomical divisions of cells in the striatum are associated with distinct gene expression specializations.

To determine whether the nine clusters corresponded to distinct cell types, we used the “self-projection method” in the Single Cell Clustering Assessment Framework (SCCAF) (Miao et al., 2020). SCCAF uses the area under the receiver operating characteristic (AUROC) curve to determine whether identified cell clusters correspond to different cell types. SCCAF split the data into a training and testing set, built classifiers based on the gene expression in the training sets, and evaluated the performance of the classifiers using the testing sets. For all nine clusters, each AUROC was more than 0.93, exceeding the recommended threshold of 0.90 for distinguishing distinct cell types (Figure 2H) (Miao et al., 2020). Moreover, the classifiers were “confused”, less than 5% of the time, even for clusters in close UMAP proximity (Figure S2D). These results indicate that the primate striatum contains at least nine distinct, region-specific cell types that all feature transcriptional characteristics of MSNs.

### MSN Type Distributions in the Dorsal Striatum

We used FISH to verify the snRNA-Seq results and explore the anatomical distribution of MSN types. To establish a baseline and to search for D1/D2-hybrids, we labeled whole coronal sections with *DRD1* and *DRD2* probes (Figure 3A). As expected, the majority of neurons expressed either *DRD1* (325 ± 99 nuclei/mm^2^) or *DRD2* (320 ± 99 nuclei/mm^2^; Figures 3B-3D). Despite the snRNA-Seq analysis suggesting that D1/D2-hybrids should constitute approximately 4% of MSNs, only ~1% of nuclei were clearly co-labeled with both *DRD1* and *DRD2* (Figure 3E). To investigate this discrepancy, adjacent sections were labeled with *DRD1*, *DRD2*, and *RXFP1*, a highly specific marker gene for the D1/D2-hybrid cluster (p = 2.47 × 10^−135^, Wilcoxon). High resolution imaging revealed that cells labeled with *DRD1* and *RXFP1* also expressed low levels of *DRD2* (Figure 3F). We used *RXFP1* and *DRD1* co-localizations to investigate the distribution of D1/D2-hybrids and found that these hybrid cells were randomly distributed in dorsal striatum (Figure 3G).

**Figure 3.**
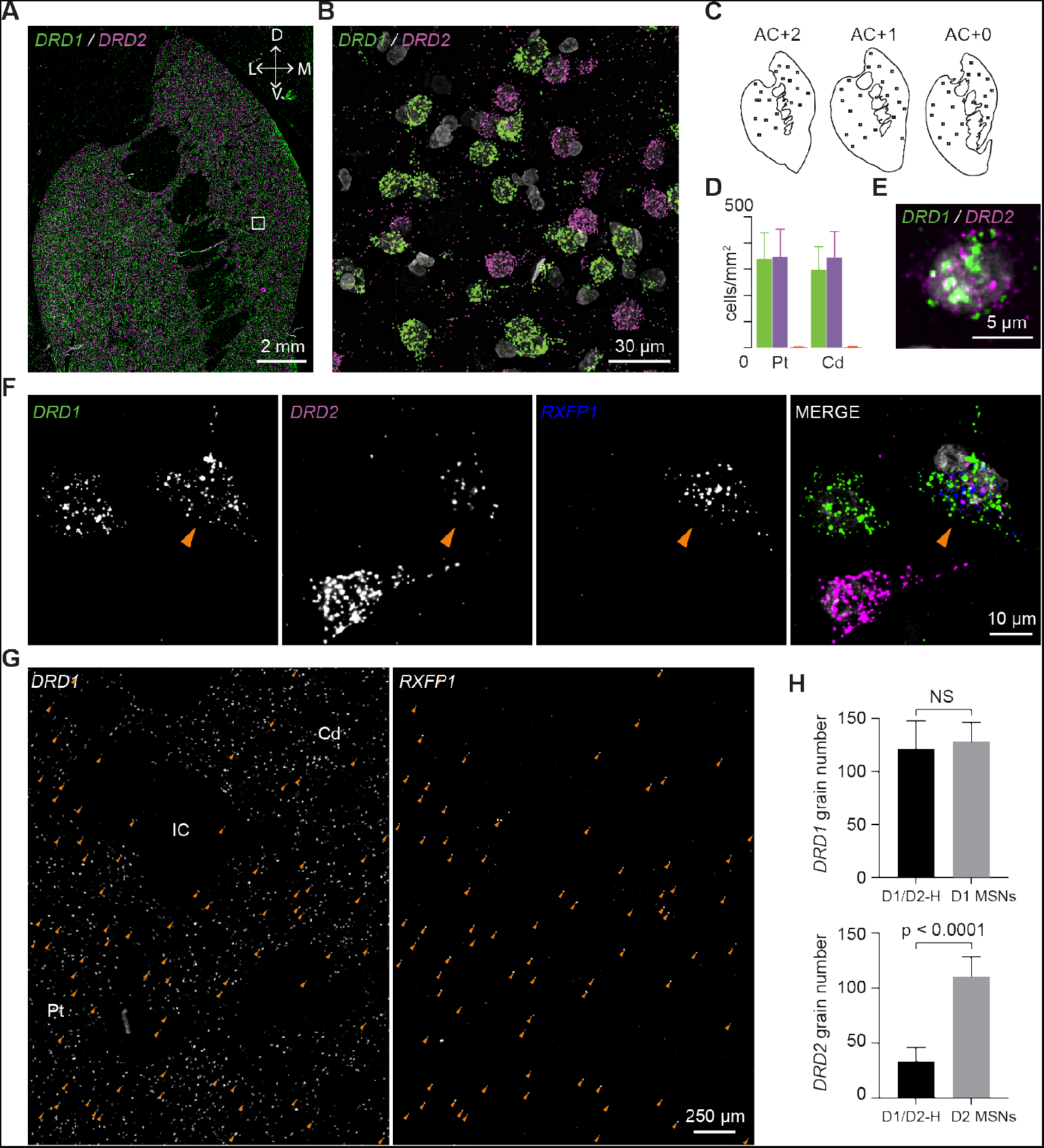
D1/D2-Hybrid Cells in Dorsal Striatum. (A)FISH labeling of *DRD1* (green) and *DRD2* (magenta). White box indicates the ROI shown in B. The top right axis shows dorsal (D), ventral (V), lateral (L), and media (M) directions. (B)High resolution image of region highlighted in A. (C)Schematic diagrams of the three sections used for *DRD1* and *DRD2* quantification. The square boxes indicate the quantified ROIs. (D)Quantification of cell density of neurons expressing *DRD1* (green), *DRD2* (magenta), or both (orange) in the caudate and putamen. Error bars are SD across ROIs. (E)One example MSN expressing both*DRD1* and *DRD2*. (F)*RXFP1* labels D1/D2-hybrid MSNs in the dorsal striatum. Arrow head points to an example D1/D2-hybrid cell. (G)FISH labeling of *DRD1* and *RXFP1*. Arrow heads point to hybrid cells. Abbreviations: Cd, caudate; Pt, putamen; IC, internal capsule. (H)Quantification of *DRD1* and *DRD2* grain number in D1/D2-hybrid cells and normal D1- or D2-MSNs. Unpaired t-test was used for statistical analysis and p values were indicated on the plots. Error bars represent standard deviation (SD) across 6 cells per type. NS: non-specific.

We compared D1/D2-hybrids to other MSN types in NHP and rodents. The number of *DRD1* grains per cell and the *DRD1* expression, measured from FISH and snRNA-Seq, respectively, in D1/D2-hybrids was not significantly different from nearby D1-MSNs (Figure 3H, p = 0.55 and 0.21 respectively, unpaired t-test). In contrast, both *DRD2* grains numbers and *DRD2* expression in D1/D2-hybrids was significantly reduced compared to nearby D2-MSNs (Figure 3H, p < 0.0001 and p = 1.17 × 10^−127^ respectively, unpaired t-test). Co-clustering of the NHP data with striatum data from recent mouse studies indicates that the D1/D2-hybrid is homologous to the recently described “D1H” (Figures S2A-S2C) (Saunders et al., 2018; Stanley et al., 2020). These results confirm the presence of a hybrid cell type in the NHP striatum.

Next, we aimed to uncover the gene signatures for the striosome and matrix compartments. Both D1- and D2-MSNs derived from the DS split into two clusters; one larger cluster and one smaller cluster. As the matrix compartment has much greater volume than the striosome compartment (Gimenez-Amaya and Graybiel, 1991; Mikula et al., 2009), we reasoned that the larger D1- and D2-MSN clusters likely corresponded to the matrix (Figure 2A, dark and light blue clusters, respectively), whereas the smaller clusters likely corresponded to striosomes (Figure 2A, light and dark red clusters, respectively). Differenital gene analysis identified marker genes for striosome and matrix compartments. The genes most enriched in the striosome D1- and D2-MSNs were *KCNT1*, *KHDRBS3*, *FAM163A*, *BACH2*, and *KCNIP1*, whereas the genes most enriched in the matrix D1- and D2-MSNs were *EPHA4, GDA*, *STXBP6*, and *SEMA3E* (Figures 2E and S3A) (p < 1.77 × 10^−52^, Wilcoxon). Further analysis revealed cell type-specific markers within the striosome compartment – *PDYN* and *POU6F2* were specifically enriched with striosome D1- and D2-MSNs, respectively (Figure S3A) (p < 1.04 × 10^−69^, Wilcoxon). We performed FISH labeling with different combinations of striosome and matrix markers. Labeling with *KCNIP1* and *STXBP6* showed the characteristic, large-scale pattern of the striosome and matrix compartments (Figures 4A and 4B), and similar results were obtained for other markers (Figures 4C, 4D and S3B-S3D). Together, these results highlight marker genes for striosome and matrix specific MSNs in the NHP DS.

**Figure 4.**
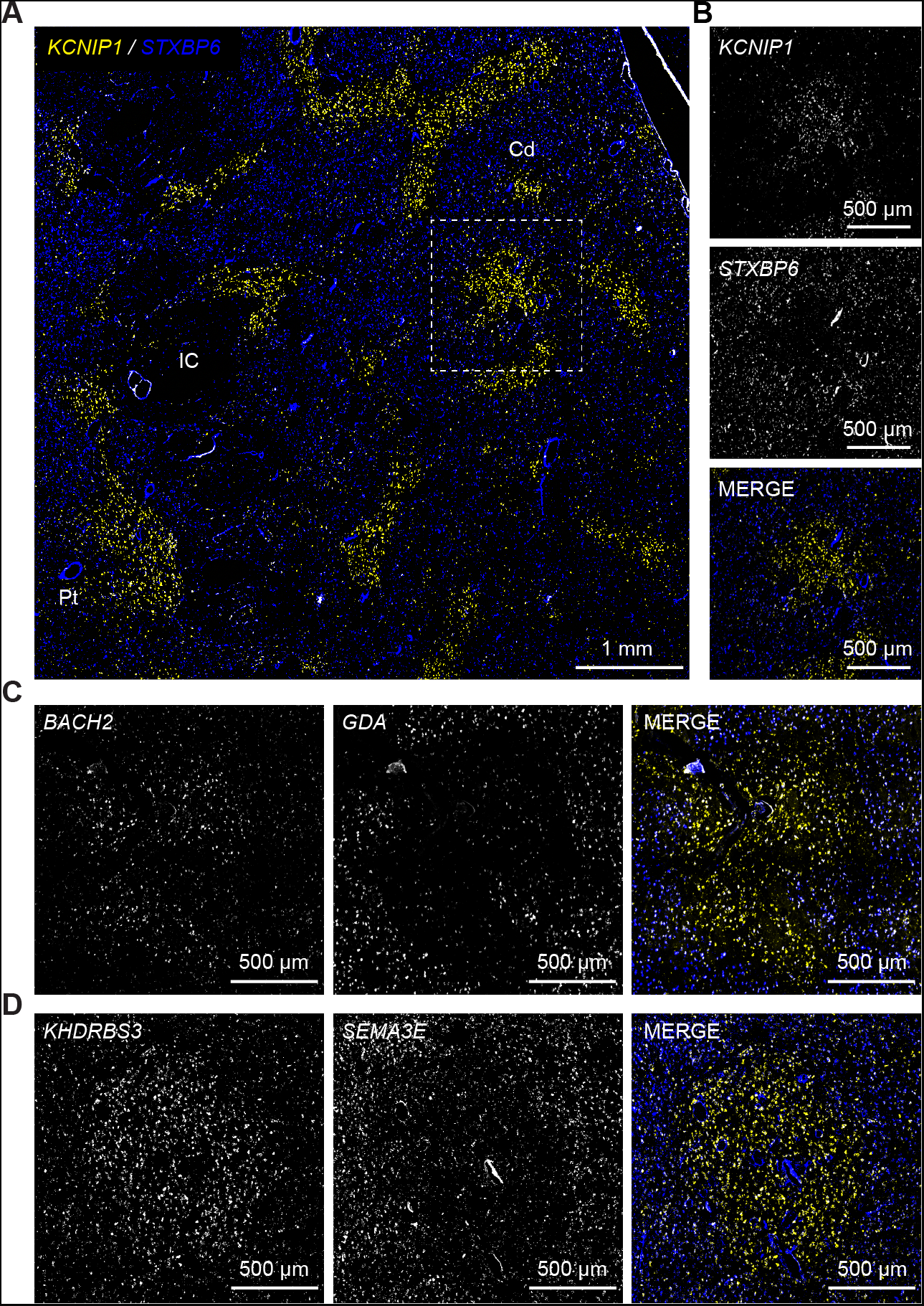
Striosome and Matrix in Dorsal Striatum. (A)FISH labeling of *KCNIP1* (yellow) and *STXBP6*(blue) distinguishes striosome and matrix, respectively. (B)Detail of the white square in F. (C)As in B, for characteristic striosome and matrix markers, *BACH2* and *GDA*. (D)As in B, for characteristic striosome and matrix markers, *KHDRBS3* and *SEMA3A*. Scale bars are indicated on images. Abbreviations: Cd, caudate; Pt, putamen; IC, internal capsule. See also Figure S3.

### MSN Type Distributions in the Ventral Striatum

Four MSN clusters were highly enriched with nuclei from the ventral striatum (Figure 2D). Three of the clusters were *DRD1* positive, whereas the fourth cluster was *DRD2* positive. Differential gene analysis of the two largest clusters (Figure 2A, light and dark green) revealed selectively enriched marker genes, including *GREB1L*, *ARHGAP6*, and *GRIA4* (Figure 2E). FISH labeling of the striatum with *DRD1* and *ARHGAP6* revealed that *ARHGAP6* was selectively enriched in a restricted portion of the ventral striatum that likely corresponded to the NAc shell and OT (Figures 5A-5C). Quantification of grain number of *ARHGAP6* in putative shell/OT and core regions confirmed that *ARHGAP6* was more enriched in shell/OT (Figure 5D). To confirm this finding, we labeled ajacent sections with *DRD1* and *GREB1L* probes and an immunofluorescent stain for calbindin. Calbindin marks the boundary between the NAc core and shell territories (Meredith et al., 1996). The *GREB1L* intensity was similar to the *ARHGAP6* intensity and traced the outline of the transition from the calbindin-poor shell to the calbinin-rich core (Figures 5F and 5G). *GREB1L* also contained more grains in shell/OT compared to core (Figure 5E). Thus, the NAc Shell and OT are comprised of region-specific D1- and D2-MSNs.

**Figure 5.**
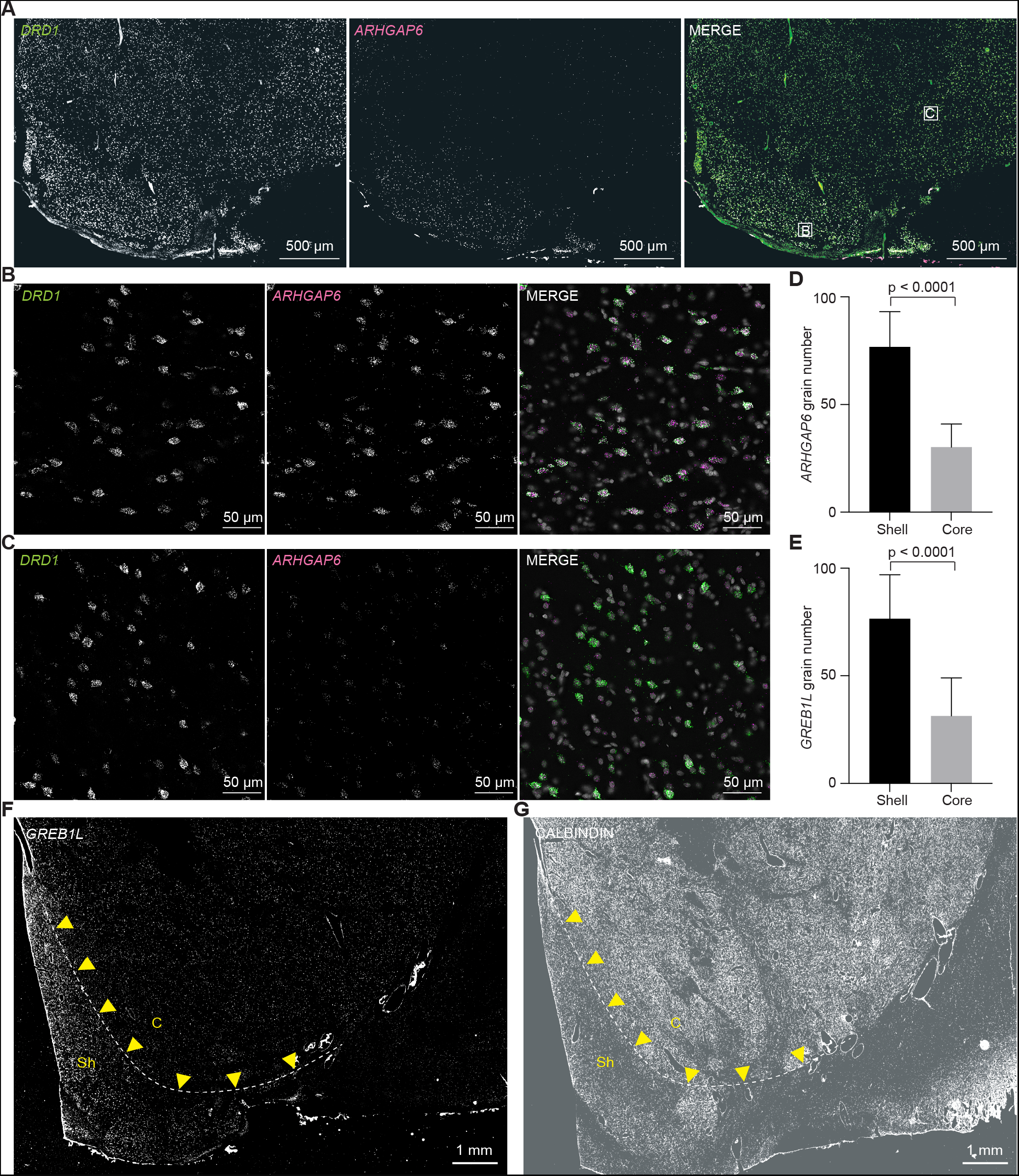
Identification of NAc Shell/OT by FISH Labeling of Gene Markers. (A)Double labeling with *DRD1* (left) and *ARHGAP6* (middle) of the NAs shell/OT shows that *ARHGAP6* is selectively enriched in the shell/OT. Lettered boxes indicate ROIs shown in B and C. (B)High-resolution image of the ROI indicated with the letter “B” in A, left. (C)As in B, for ROI indicated with letter “C”.(D)Quantification of grain number of *ARHGAP6* in shell/OT and core. Unpaired t-test was used for statistical analysis. Error bars represent standard deviation (SD) across 32 cells per each region. (E)Quantification of grain number of *GREB1L* in shell/OT and core. Unpaired t-test was used for statistical analysis. Error bars represent standard deviation (SD) across 29 cells per each region. (F)FISH labelling of *GREB1L* and *GREB1L* intensity follows the border of the shell (dashed white line), indicated by the yellow arrows. (G)Calbindin immunohistochemistry reveals the border between the core and shell (dashed white line), highlighted by yellow arrows.

One of the most remarkable features of the *DRD1* and *DRD2* labelling in the VS was the presence of D1-exclusive islands that likely corresponded to “interface islands” (Figure 6A) (Heimer, 2000; Prensa et al., 2003). We investigated whether the cell types within these D1-exclusive islands corresponded to the smaller*DRD1* enriched VS clusters (Figure 2A, light and dark orange). We selected two respective marker genes, *RXFP1* and *CPNE4*, for the two *DRD1* enriched VS clusters (Figure S5C). We labeled one section using *DRD1*, *RXFP1,* and *CPNE4* probes and revealed that *RXFP1* and *CPNE4* labeled distinct D1-exclusive cell islands (Figure 6B). High resolution confocal microscopy of these islands verified that *CPNE4* and *DRD1* colocalized in the same cells in one island (Figure 6C), whereas *RXFP1* and *DRD1* colocalized in the same cells of another island (Figure 6D). These results suggest that different interface islands contain different *DRD1*-positive cell types.

**Figure 6.**
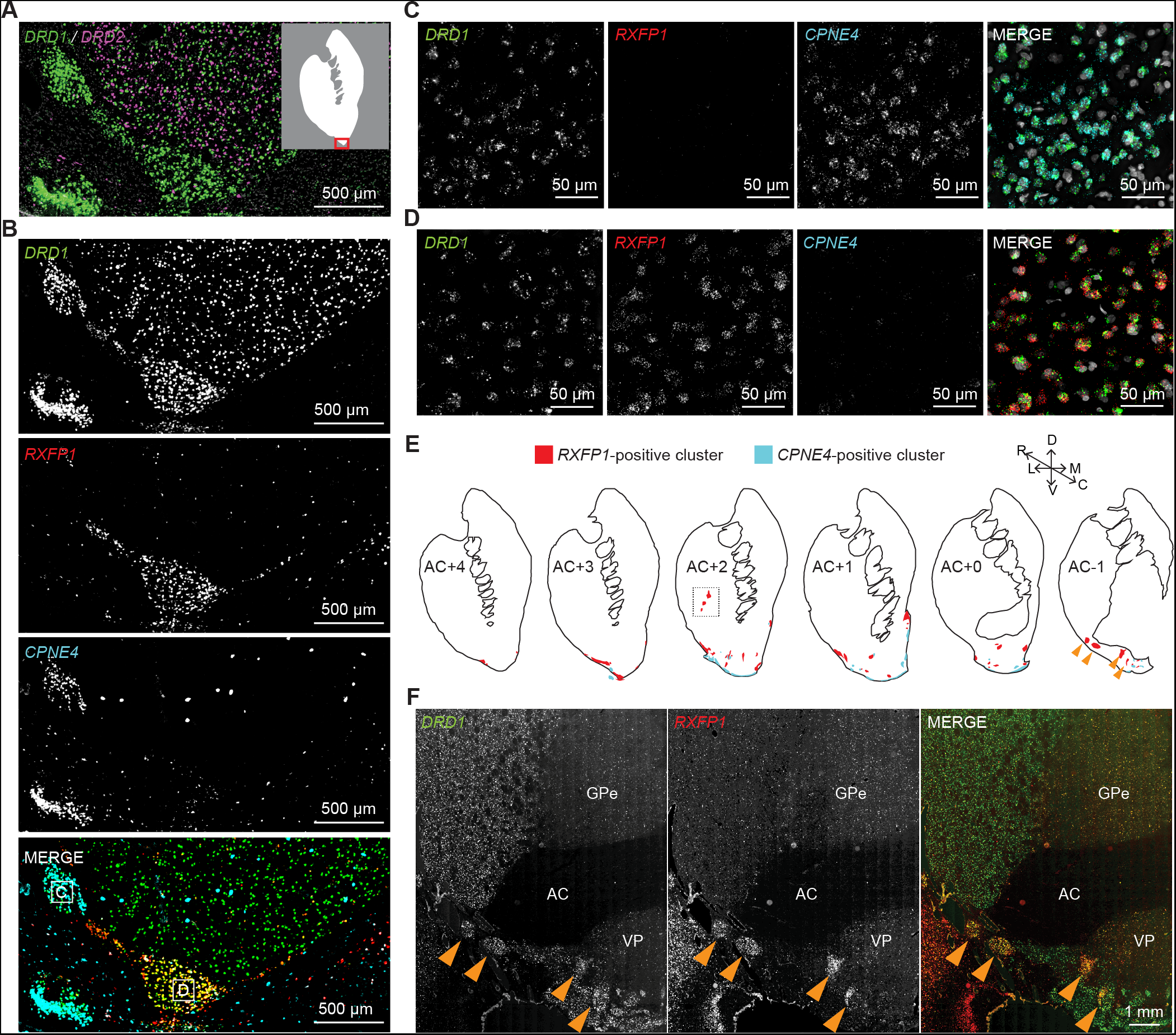
Identification and Distribution of Two Interface Islands in VS. (A)FISH stain of *DRD1* (green) and *DRD2* (magenta). *Inset*: White area indicates striatum and the red box highlights the area shown in A and B. (B)FISH stain of *DRD1*, *RXFP1* and *CPNE4* in immediately adjacent section from A. (C)High-resolution image of the ROI indicated with the letter “C” in B. (D)High-resolution image of the ROI indicated with the letter “D” in B. (E)Distribution of *RXFP1* and *CPNE4* clusters across six rostral-caudal regions identified by multichannel FISH. The upper right axis shows dorsal (D), ventral (V), lateral (L), medial (M), rostral (R), and caudal (C) directions. Dashed white box denotes *RXFP1* clusters in putamen, the FISH image of which is shown in Figure S5H. Yellow arrowheads point to *RXFP1* clusters in caudal extent of NAc in both illustration and images shown in F. (F)Example *RXFP1* clusters in caudal extent of NAc. AC = anterior commissure, GPe = external globus pallidus, VP = ventral pallidum. See also Figure S4 and S5.

To identify the interface islands and determine their distribution, we repeated the *DRD1*, *RXFP1*, and *CPNE4* FISH on regularly spaced pre- and peri-commissural coronal sections. We defined the regions of the VS by comparison of Nissl-stained sections with a high-resolution MRI and DTI Rhesus macaque brain atlas (Figures S5A and S5B) (Bakker et al., 2015; Calabrese et al., 2015; Paxinos et al., 2000) and we mapped all the nearby islands in two monkeys (Figures 6E and S4). The *CPNE4*-positive islands appeared to correspond to the Islands of Calleja (ICj) (Meyer et al., 1989). This correspondence was verified by intense co-localization of *CPNE4* and *DRD1* in cells in the major ICj, an easily identifiable landmark at the border between the NAc and the septal nuclei (Figure S5D). Previous studies have shown that cells in ICjs are granule cells (Meyer et al., 1989). Likewise, the *CPNE4*-positive cells we examined were small and exhibited high packing density (Figures S5E and S5F). Thus, the cluster enriched with *DRD1* and *CPNE4* corresponded with granule cells in the ICjs. Outside of the ICjs, co-localization between *DRD1* and *CPNE4* was restricted to a dense cell layer at the ventral extreme of the OT, possibly corresponding to a portion of the anterior olfactory nucleus (AON) (Figures 6B and S5G). Gene enrichment analysis revealed that differentially expressed genes in these cells are implicated in neurogenesis, neurosecretion, and many other functions (Table S3).

In contrast to the *CPNE4*-positive ICjs, the *RXFP1-*positive islands had larger nuclei that were less densely packed together and did not appear different from nearby D1-MSNs (Figures S5E and S5F). Likewise, *RXFP1*-positive islands were not restricted to the border regions of the VS, rather, they were found throughout the NAc, putamen, and near the adjacent septal nuclei (Figure 6E, orange arrows and dashed black box, Figures S4, S5H, S5I, and S6A). *RXFP1*-positive cells located in these VS islands exhibited high levels of *DRD1* expression, but no detectable *DRD2* expression (Figures S6A and S6B). Moreover, cells in the *RXFP1*-positive islands expressed high levels of the gene for the μ-opioid receptor (*OPRM1*) compared to D1-MSNs located nearby but outside of the D1-exclusive island (Figures 7A and 7B). Previous research using μ-opioid receptor (MOR) radioligand binding revealed MOR-rich islands in the NAc, putamen, and nearby basal forebrain and designated them as Neurochemically Unique Domains in the Accumbens and Putamen (NUDAPs) (Daunais et al., 2001; Voorn et al., 1996). Based on their distribution and the upregulation of *OPRM1* – upregulation that was not present in the ICjs (Figures 7C and 7D) – we concluded that these *RXFP1*-positive interface islands corresponded with NUDAPs, and we denoted the *DRD1*- and *RXFP1*-positive cells as D1-NUDAP neurons. Gene enrichment analysis revealed that D1-NUDAP neurons express genes that have been implicated in drug addiction and many other functions (Table S3). These data show a novel cell type that is associated with particular interface islands and that could be critical for the hedonic aspects of reward. Altogether, these results demonstrate that the ventral striatum is characterized by the presence of discrete cell types that correspond to functionally relevant subdivisions including NAc shell, OT, and distinct types of interface islands.

**Figure 7.**
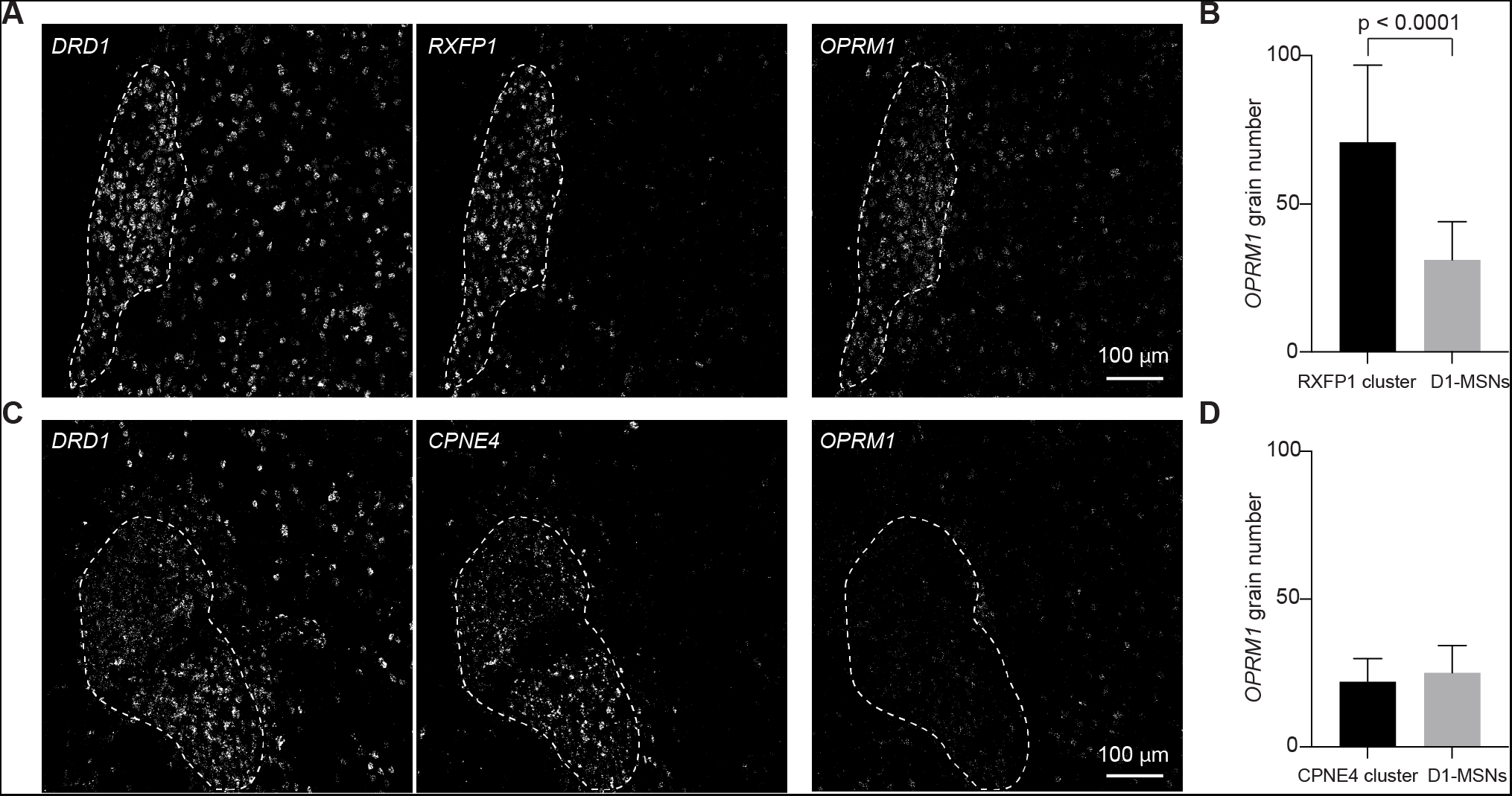
*RXFP1* Clusters Corresponds to NUDAPs. (A)FISH stain of *DRD1* and *RXFP1* as well as *OPRM1* in two close sections. White dashed line delineates the boundaries of a *RXFP1* cluster. (B)Quantification of grain number of *OPRM1* in *RXFP1* clusters and close D1-MSNs. Unpaired t-test was used for statistical analysis. Error bars represent standard deviation (SD) across 41 cells in each group. (C)FISH stain of *DRD1* and *CPNE4* as well as *OPRM1* in two close sections. White dashed line delineates the boundaries of a *CPNE4* cluster. (D)Quantification of grain number of *OPRM1* in *CPNE4* clusters and close D1-MSNs. Unpaired t-test was used for statistical analysis. Error bars represent standard deviation (SD) across 36 cells in each group. See also Figure S6 and S7.

## DISCUSSION

Here we investigated the transcriptional and anatomical diversity among medium spiny neurons (MSNs) and MSN-like neurons in the non-human primate (NHP). Single nucleus RNA-sequencing (snRNA-Seq) revealed a clear distinction between MSN types located in the dorsal and ventral striatum. The dorsal striatum included D1- and D2-MSN types specific to the striosome and matrix compartments (Figure 2A). We found marker genes, including *BACH2* and *KCNT1* (Figures 2E and 4), that were expressed in all striosome MSNs, but we also found markers for compartment-specific cell types, including *PDYN* and *POU6F2* that were enriched in D1- and D2-striosome MSNs, respectively. The ventral striatum also yielded distinct MSN clusters. Two of the clusters corresponded to D1- and D2-MSNs located in the NAc shell and OT (Figures 2A and 5). Transcriptional and FISH analysis of the other two clusters revealed them to be associated with the “interface islands” – dense cellular islands located within and near the ventral border of the striatum. One of these clusters was highly enriched with *CPNE4* and strongly labeled the ICjs and select cells in the OT. The VS D1-MSN cluster marked by expression of *RXFP1* mapped to NUDAPs, regions previously identified in human and NHP nucleus accumbens and putamen via μ-opioid receptor ligand binding (Daunais et al., 2001; Voorn et al., 1996). Together, these cell types provide a blueprint for studying cell type-specific functions during sophisticated primate behaviors, and the cell type-specific marker genes define potential molecular access points to enable the application of genetically coded tools in scientific or translational contexts.

The dorsal striatum samples revealed D1- and D2-MSN types associated with striosome and matrix compartments, as well as a fifth type that was a *DRD1* and *DRD2* hybrid (D1/D2-hybrid). Statistical examination of gene expression levels confirmed that members of this hybrid class were not doublets. Comparison of the D1/D2-hybrid with rodent single cell studies revealed these neurons to be a novel class of MSNs recently described by several groups and referred to as “D1H” and “eccentric SPNs” (Martin et al., 2019; Saunders et al., 2018; Stanley et al., 2020). Specifically, co-clustering of the NHP data with the data from Stanley et al., 2020, showed that approximately half of the “D1H” population co-clustered with the D1/D2-hybrid. Likewise, the D1/D2-hybrid expressed many genes that are enriched in D1H and eSPNs, including *FOXP2* and *CASZ1* (Martin et al., 2019; Saunders et al., 2018; Stanley et al., 2020). FISH analysis of the D1/D2-hybrid cells revealed a spatial distribution similar to eSPN distribution (Figure 3G) (Saunders et al., 2018). Interestingly, these cells were enriched with gene markers that we and others have found in the striosome compartment, including *BACH2*, *KCNT1*, *OPRM1*, *KCNIP1*, and *KHDRBS3* (Figure S6C), but the FISH analysis did not show that these hybrids were preferentially located in striosome (Figure 3G). These results lead us to wonder whether there is a relationship between D1/D2-hybrid cells and “exo-patch” cells: striosome-like neurons scattered throughout the matrix (Smith et al., 2016). Future studies will be required to explore this link, but these results indicate that this novel cell type is conserved in the primate lineage.

The ventral striatum samples revealed a subpopulation of MSN-like cells that we designated as D1- NUDAP neurons, because of their resemblance to dense patches of μ-opioid receptor ligand binding in the ventral striatum named Neurochemically Unique Domains in the Accumbens and Putamen (NUDAPs) (Daunais et al., 2001; Voorn et al., 1996). D1-NUDAP neurons shared some characteristics with D1/D2-hybrid neurons. For instance, both cell types expressed *RXFP1* as a marker gene and both were enriched with *OPRM1*. Likewise, co-clustering with mouse single cell data revealed that approximately half of the D1H population co-clustered with D1/D2-hybrids, and the other half co-clustered with D1-NUDAPs (Figures S2A-S2C). Despite these similarities, our data indicate that D1-NUDAP neurons and D1/D2-hybrid neurons represent distinct cell types. First, although *DRD2* expression was common in D1/D2-hybrids (Figures 2D and 3F), we found no evidence of *DRD2* expression in D1-NUDAPs (Figure S6). Second, the D1-NUDAP neurons did not express marker genes, including *CASZ1* and *GRIK1,* that were reported in D1/D2-hybrids, D1H, and eSPNs, *(Martin et al., 2019; Saunders et al., 2018; Stanley et al., 2020)*. Finally, a machine learning classifier easily distinguished between D1-NUDAP and D1/D2-hybrid neuron types (Figure 2H). Thus, these data reveal cell type-specific interface islands, NUDAPs, that are composed of a *DRD1*-positive cell type not found elsewhere in the striatum.

The functions of the interface islands – both ICjs and NUDAPs – remains mysterious. ICjs are embedded within the border region of the olfactory tubercle, and it has therefore been reasoned that they play a role in olfactory processing. However, a comparative study across mammal species revealed that the size of the major island, proportional to the rest of the brain, reached its greatest extent in man, a microsmatic creature, rather than in macrosmatic creatures with highly developed senses of smell, including mice, rats, and hedgehogs (Meyer et al., 1989). Moreover, dolphins, who completely lack a sense of smell, possess well developed ICjs. These observations are not consistent with the hypothesis that primary function of the ICjs is olfactory processing. An alternative role for ICjs has been proposed as a pool of immature neurons from which adult striatal neurogenesis is derived (Ernst et al., 2014). Several lines of evidence are consistent with this hypothesis. A longitudinal study quantified the size of the major ICj across the life span of cats and showed that its size decreased with age (Meyer et al., 1989), as might be expected from pools of immature neurons that are slowly drained. This notion is further backed up by studies that have shown the ICjs are enriched with *BCL2* – a marker gene for immature neurons (Bernier and Parent, 1998; Fudge and Haber, 2002). Indeed, our snRNA-Seq data and FISH analysis showed that both the D1-ICjs and the D1-NUDAPs are strongly enriched with *BCL2* (Figures S5C and S7). Finally, our gene enrichment analysis was reasonably conclusive with a role for ICjs in adult neurogenesis (Table S3). Another enticing possibility is that the interface islands have a neurosecretory role. The islands are consistently observed near blood vessels (Meyer et al., 1989; Prensa et al., 2003) and our gene enrichment analysis indicates that both the D1-ICjs and the D1-NUDAPs were enriched with genes implicated in neurosecretory functions (Table S3). In fact, one of the best marker genes for the D1-NUDAP neurons is *RXFP1* – a member of the relaxin receptor family that is broadly implicated in neuroendocrine function. Likewise, *CPNE4*, the marker for the D1-ICjs, might participate in secretion through promoting membrane fusion (Koukalova et al., 2018). There is a tight link between limbic processing and autonomic responses to emotional stimuli. A neurosecretory role for the interface islands would present a novel pathway for the brain to affect the body and support this link. In either scenario, our data show that ICjs and NUDAPs are comprised of distinct cell types. The cell type-specific gene expression data can highlight molecular access points, and these access points will enable the use of genetically coded tools to discover the functional properties of the interface islands and their constituent neurons.

Both the striosome compartment in the DS and the cell types in the VS are implicated in limbic and reward processing. It is therefore interesting to speculate on the relationship between the striosome compartment and the VS cell types. Our data indicate that there are many transcriptional similarities between the striosome and the NUDAPs, but also some key differences. Many striosome specific markers are upregulated in NUDAPs, including *KCNIP1*, *KCNT1*, *KHDRBS3*, and *BACH2* (Figure S6C). Even *PDYN*, which is a widely acknowledged D1-striosome marker gene (Xiao et al., 2020), is also expressed in D1-NUDAPs. On the other hand, D1-NUDAP cells also express some genes which we found to be selectively expressed in the matrix, including *STXBP6*, *GDA*, and *SEMA3E*. Finally, it is worth discussing *OPRM1*, the gene for the μ-opioid receptor. Many studies use *OPRM1* as a marker for striosomes (Märtin et al., 2019). Our data revealed that *OPRM1* was upregulated in the striosome, as predicted, but the upregulation was not as dramatic as we expected. In contrast, we observed strong *OPRM1* signals in the NUDAPs (Figures 7A and 7B). Indeed, this selective enrichment in *RXFP1*-positive interface islands is a key piece of evidence in favor of the NUDAP hypothesis. This selective *OPRM1* enrichment suggests another tantalizing link: could NUDAPs be part of the network of “hedonic hotspots?” Hedonic hotspots are regions in the NAc and ventral pallidum that, when opioids are directly applied, produce behavioral reactions that indicate pleasure (Berridge and Kringelbach, 2015). The differentially expressed genes presented here provide a blueprint for answering this question and studying other cell type-specific contributions to reward processing.

The data we show here indicates that there are discrete cell types in well-defined subregions of the striatum. At first, this conclusion might seem to be at odds with findings from rodents that illustrate continuous variation (Stanley et al., 2020). However, we do not think there is any conflict between these studies. We selected FISH markers based on their ability to discriminate cell types, and in doing so we conclusively show that clear demarcations are prevalent throughout the striatum. Nonetheless, there are likely many other marker genes that would show continuous variation in expression levels, and we are confident that a more extensive search would reveal more features, both discrete and continuous. Indeed, a limitation of this work was our inability to isolate and identify MSN cell types associated with the core regions of the NAc. We found several gene markers, including *ARHGAP6*, *GREB1L*, and *GRIA4* that were upregulated in the VS samples (Figure 2). FISH labeling with these probes revealed that they were expressed in the NAc shell and OT (Figure 5). In fact, the FISH signals traced the transition of Calbindin poor shell to the Calbindin rich core. Thus, these markers highlighted the ventral border of the core, but we did not find markers that separated the core from the DS. There was a significant fraction of nuclei that were derived from the VS sample, but that clustered with D1- and D2-matrix cell types (Figure S1M). The border between the NAc core and the DS is not clearly demarcated, and a likely explanation for this clustering result is that we inadvertently included the ventral DS in our VS samples. However, this does not explain the absence of a clear core signal. Explaining this will require careful future experiments. Given the substantial literature showing NAc subdivisions and their diverse roles in reward processing, decision making and drug addiction (Calipari et al., 2016; Castro and Bruchas, 2019; Floresco, 2015; Hyman et al., 2006), further experiments to unambiguously sample these territories and define their component cell types is strongly warranted.

This and other studies demonstrate that single cell technologies can be effectively applied to NHPs (Krienen et al., 2020; Zhu et al., 2018). We foresee at least three critical reasons to avidly apply single cell technologies to NHPs. (1) Old-world monkeys like Rhesus macaques are more similar to humans than any other research animal that allows for invasive neurophysiological experiments. Accordingly, Rhesus macaque cell types, including highly specialized neurons like Betz cells, von Economo neurons, and even striatal interneuron types, recapitulate homologous human cell types better than cells from rodents or even from marmosets (Bakken et al., 2020; Evrard et al., 2012; Krienen et al., 2020). (2) Single cell studies performed on post-mortem human tissue are subject to different ethical constraints that manifest as relatively long and highly variable postmortem intervals (Hodge et al., 2019). In contrast, NHP experiments can be performed in a highly controlled and more timely fashion. (3) NHPs have resisted the widespread application of modern genetically coded tools. Single cell technologies, including RNA-Seq, ATAC-Seq, and bisulfide-seq can identify atomized cell types and potent regulatory sequences that will break this resistance and enable the effective applications of genetically coded tools in large, wild type animals that resemble humans. Rhesus macaque behavior is readily interpretable in terms of human behavioral theory such as economic theory (Genest et al., 2016; Lak et al., 2014; Stauffer et al., 2015; Stauffer et al., 2014), learning theory (Waelti et al., 2001), and even game theory (Barraclough et al., 2004; Haroush and Williams, 2015). The diversity of MSN cell types presented here provides a blueprint to investigate the cell type-specific mechanisms for such sophisticated behaviors.

## ACKNOWLEDGMENTS

We would like to thank Andreea Bostan for constructive and thoughtful discussions. We thank Jacquelyn Breter for taking excellent care of animals. We also thank Jaimi Nagashima and Mitsutoshi Hanada for their assistance with large image scanning. This works was supported by NIH grants UG3MH120094 (WRS) and DP2MH113095 (WRS).

## AUTHOR CONTRIBUTIONS

JH, LCB, ARP, and WRS designed the research project. JH, LCB, AA, KMR, BEO, MW, KF, and WRS collected the data. JH, MK, JC, ARP, and WRS analyzed the data. JH and WRS wrote the paper.

## CONTACT FOR REAGENT AND RESOURCE SHARING

Further information and requests for reagents and resources should be directed to and will be fulfilled by the Lead Contact, William R. Stauffer.

## EXPERIMENTAL MODEL AND SUBJECT DETAILS

### NHPs

All animal procedures were in accordance with the National Institutes of Health Guide for the Care and Use of Laboratory Animals and approved by the University of Pittsburgh’s Institutional Animal Care and Use Committee (IACUC) (Protocol ID, 19024431). Rhesus Monkeys were single- or pair-housed with a 12h-12h light-dark cycle. Monkey F was a 12-year-old female (8.1 kg). Monkey P was a 5-year-old female (5.4kg). Monkey B was a 13-year-old male (11.78 kg). Monkey K was a 4-year-old male (6.0 kg).

## METHOD DETAILS

### MRI and Surgery

For MRI, we anesthetized monkey F and P with ketamine and maintained general anesthesia with isoflurane. We head fixed the monkeys using a MRI-compatible stereotaxic frame and scanned (Siemens, 3T) for anatomical MRI. We generated a whole brain model for each monkey using Brainsight (Rogue Research) and 3-D printed a custom matrix for the brain with cutting guide set every 1 mm.

To maximize nuclei viability, we followed a harvesting protocol similar to the one outlined in Davenport et al. (2014). Briefly, animals were initially sedated with ketamine (15 mg/kg IM), and then ventilated and further anesthetized with isoflurane. The animals were transported to a surgery suite and placed in a stereotaxic frame (Kopf Instruments). We removed the calvarium and then perfused the circulatory systems with 3-4 liters of ice cold artificial cerebrospinal fluid (ACSF; 124 mM NaCl, 5 mM KCl, 2 mM MgSO_4_, 2 mM CaCl_2_, 23 mM NaHCO_3_, 3 mM NaH_2_PO_4_, 10 mM glucose; pH 7.4, osmolarity 290–300 mOsm) oxygenated with 95 % O_2_:5 % CO_2_. We then opened the dura and removed the brain. We sliced on the custom brain matrix into 4 mm slabs. We removed three striatal regions – the caudate, putamen, and ventral striatum –under a dissection microscope for nuclei isolation. Monkeys B and K, for FISH, were perfused with 4% paraformaldehyde (PFA, Sigma-Aldrich, Cat# P6148) in phosphate buffered saline (PBS, Fisher scientific, Cat# BP243820) supplemented with 10% sucrose (Sigma-Aldrich, Cat# S8501). The brain was post-fixed with 4% PFA and cryopreserved with a gradient of sucrose (10%, 20%, 30%) in PBS.

### Nuclei isolation

We isolated nuclei isolated as previously described (Habib et al., 2017). Briefly, we homogenized tissues using a loose glass dounce homogenizer followed by a tight glass homogenizer in EZ PREP buffer (Millipore Sigma, Cat# NUC-101). We washed nuclei once with EZ PREP buffer and once with Nuclei Suspension Buffer (NSB; consisting of 1× PBS, 0.01% BSA and 0.1% RNase inhibitor (Clontech, Cat# 2313A)). We re-suspended the washed nuclei in NSB and filtered them through a 35-μm cell strainer (Corning, Cat# 352235). We counted the nuclei and diluted down to 1000 cells/μl. We loaded approximately 10,000 cells from each brain region onto a 10X chip which were then run through a 10x Genomics Chromium controller.

### Single nucleus RNA-Seq

We used 10x Chromium Single Cell 3’ Reagent kits v3 Chemistry (10x Genomics, Cat# PN-1000075) for monkeys F and P. We reverse transcribed RNAs and generated libraries according to 10x Genomics protocol. Briefly, we generated Gel beads-in-emulsion (GEMs) after running through a 10x Genomics Chromium controller. We reverse transcribed mRNAs within GEMs in a Bio-Rad PCR machine (Cat# C1000). We barcoded cDNAs from individual cells with 10x Genomics Barcodes and barcoded different transcripts with unique molecular identifiers (UMIs). We purified cDNAs with Dynabeads (10x Genomics, Cat# 2000048) after breaking the emulsion with a recovery agent (10x Genomics, 220016). Then, we amplified cDNAs by PCR and purified them with SPRIselect reagent (Beckman Coulter, Cat# B23318). We analyzed the cDNA quantification and quality using Agilent Bioanalyzer 2100. We prepared libraries following fragmentation, end repair, A-tailing, adaptor ligation, and sample index PCR. We quantified the libraries by qPCR using a KAPA Library Quantification Kit (KAPA Biosystems, Cat# KK4824). We pooled together libraries from individual monkeys and loaded them onto NovaSeq S4 Flow Cell Chip. We sequenced samples from monkeys F and P to depths of 400,000 and 250,000 reads per nuclei, respectively.

### Immunohistochemistry for Calbindin

We sought to verify previously labeled sections containing FISH probes for possible shell markers using adjacent tissue sections and immunohistochemistry to label for Calbindin. The sections were rinsed in Phosphate Tris (PT, pH 7.2-7.4) buffer and blocked in 10% normal donkey serum (Jackson, Cat# 017-000-121) solution for one hour. The sections were then incubated with primary antibody solution (Swant, Calbindin D- 28K, 300, 1: 7,000) overnight at 4°C. The sections were then rinsed in PT buffer and incubated in secondary solution (Alexa Fluor 568, Donkey anti-mouse, 1:300) at room temperature for two hours. They were then rinsed in PT buffer and counterstained with Hoechst (Invitrogen, Cat# H21486, 1: 10,000) for 10 minutes. Lastly, the sections were rinsed in PT buffer and mounted with Prolong Gold Antifade (ThermoFisher, Cat# P36930).

### FISH probes

We ordered custom FISH probes from ACD to validate direct- and indirect- pathways and NAc-enriched MSN cell types, as follows: *DRD1* (ACD #549041, a 20ZZ probe targeting 1335-2279 of NM_001206975.1), *DRD2* (ACD #549031-C2, a 20ZZ probe targeting 232-1470 of XM_001085571.3), *RXFP1* (ACD #801121-C2, a 20ZZ probe targeting 1508-2592 of XM_001096574.4), *CPNE4* (ACD #801111-C3, a 20ZZ probe targeting of 2-943 of XM_028843706.1), *KCNIP1* (ACD #889143-C3, a 20ZZ probe targeting 283-1626 of XM_015141392.2), *KCNT1* (ACD #881571-C3, a 20ZZ probe targeting 964-1906 of XM_015116381.2), *BACH2* (ACD #898961-C3, a 20ZZ probe targeting 927-2260 of XM_028847343.1), *KHDRBS3* (ACD #881591-C3, a 20ZZ probe targeting 494-1493 of XM_028852934.1), *STXBP6* (ACD #881611-C2, a 20ZZ probe targeting 2-1077 of NM_001260925.1), *SEMA3E* (ACD #879971-C2, a 20ZZ probe targeting 905-1889 of XM_028846126.1), *GDA* (ACD #881601-C2, a 20ZZ probe targeting 355-1373 of XM_015117899.2), *GREB1L* (ACD #898991-C3, a 20ZZ probe targeting 784-1732 of XM_015121640.2), *ARHGAP6* (ACD #898981-C3, a 20ZZ probe targeting 2516-3463 of XM_001094565.3).

### FISH stain and imaging

We embedded brains in optimal cutting temperature (OCT) and stored them at −80°C until cutting. We cut floating sections at 15 and 30 μm, in monkey B and K, respectively, mounted tissue on 2×3” pre-treated slides, and preserved the slides in a freezer at −80°C. We used the Advanced Cell Diagnostics (ACD) RNAscope platform and Multiplex Fluorescent Detection Reagents v2 (ACD, Cat# 323110) to perform FISH with slight modifications for monkey brain tissue. We air dried slides for 30 min after removal from −80°C freezer and baked them at 60°C for 20 min. We treated the brain sections with hydrogen peroxide (ACD, Cat# 322335) for 10 min at room temperature and then with RNAscope Target Retrieval Reagents (ACD, Cat# 322000) for 8 min at 99°C. We incubated the slides in 100% alcohol for 3 min and then dried them in 60°C for 10 min. We treated the samples with protease III (ACD, Cat# 322337) for 30 min at 40°C and incubated them with probes for 2 hr. After hybridization with AMP 1, 2 and 3, we incubated the slides with different HRP channels and fluorophores, including Opal 520 (PerkinElmer, Cat# FP1487A), Opal 570 (PerkinElmer, Cat# FP1488A) and Opal 650 (PerkinElmer, Cat# FP1496A). Lastly, we used Trueblack (Biotium, Cat# 23007) to quench Lipofuscin autofluorescence for 45 s at room temperature and counterstained every slide with DAPI before mounting with Prolong Gold Antifade Mountant (Life technologies, Cat# P36930).

We scanned labeled sections using a Hamamatsu NanoZoomer S360 or a Nikon Eclipse T*i*2 under 20× objective. For high resolution images, we used a Nikon Eclipse T*i*2 or an Olympus IX83 under 40× or 63× objective or 40× with additional 2× built in objective (equal to 80×). We took multi-layer images and did deconvolution using Nikon’s NIS-Elements deconvolution software and used maximum intensity projections to create single images. We used NDP software to convert the NanoZoomer original files to tiff format and ImageJ and Adobe Photoshop to adjust brightness and overlay images.

## QUANTIFICATION AND STATISTICAL ANALYSIS

### Custom annotation file

We downloaded the macaque rheMac10 genome (Warren et al., 2020) (https://hgdownload.soe.ucsc.edu/goldenPath/rheMac10/bigZips/rheMac10.fa.gz) and human NCBI RefSeq transcriptome annotation gtf file from the UCSC genome browser (https://hgdownload.soe.ucsc.edu/goldenPath/hg38/bigZips/genes/hg38.ncbiRefSeq.gtf.gz). We used the UCSC liftOver tool with the hg38toRheMac10 chain file (https://hgdownload.soe.ucsc.edu/goldenPath/hg38/liftOver/hg38ToRheMac10.over.chain.gz) to overlay the human transcriptome gtf file onto the rheMac10 genome, with a minimal match threshold of 0.85. The human “liftOver” annotations were used to extend and supplement the macaque annotations, leading to greater numbers of genes called (Figure S1A).

### Data analysis

#### Preliminary exploratory analysis

We performed exploratory analysis of striatum samples in monkey F. We converted raw illumina signal bcl to fastq files using the Cell Ranger suite, then aligned to the rheMac10 genome using the STAR aligner with the - solo option for single cell demultiplexing and using the custom rheMac10 gtf as above. This yielded 3,751,710 nuclei. We removed empty droplets with the dropletUtils *defaultDrops* function (Griffiths et al., 2018), and removed doublets using the *SCDS* hybrid doublet caller with a threshold of 1.0 (Bais and Kostka, 2020), lowering the number of nuclei to 8,700. We performed pre-clustering using Scanpy’s Leiden clustering function with low quality cells removed based on gene count with cluster-specific parameters (Wolf et al., 2018), to bring the total number of nuclei in our analysis to 6,199. A total of 34,722 genes were identified, with an average of 4,618 per nucleus. We normalized single cell gene expression values using the *scran* size factor normalization method, with our pre-clustering as input to the function (Lun et al., 2016). After expression count normalization, we used Scanpy’s highly variable genes function to select the top 7500 highly variable genes in the dataset. We performed an iterative principal component analysis (PCA) using the highly variable genes from nearest neighbor graphs with Scanpy’s *neighbors* function. We used Leiden clustering of the resulting neighbor graphs to distinguish major cell types (MSN, interneurons, astrocytes, microglia, oligodendrocytes, oligodendrocyte precursors, endothelial cells, and mural/fibroblast-like cells). We performed Leiden clustering on medium spiny neurons specifically, resulting in a D2 neuron cluster and several robust D1 neuronal types. The relevant code is available at https://github.com/pfenninglab/nhp_snrna_striatum_analysis.

#### Data integration analysis

To expand the scope of the paper to cover both dorsal and ventral striatum, and to replicate the findings from monkey F, we collected another high-quality data set from a new monkey P. Alignment of the reads by cellranger count to the rheMac10 genome using the custom rheMac10 gtf yielded 61,609 and 23,690 nuclei for monkey P and F respectively. We used a Seurat v3 pipeline to perform an integrated analysis of monkey F and P (85,299 nuclei in total). First, we removed ambient RNA in monkey P using SoupX (Young and Behjati, 2020). We then deleted ribosomal genes for both monkeys, and removed doublets using DoubletDetection (McGinnis et al., 2019), which lowered the number of nuclei to 80,902. Neuronal cells express higher numbers of genes compared to non-neuronal cells (Hodge et al., 2019; Tasic et al., 2018; Zeisel et al., 2018) and therefore we used two different thresholds to remove low quality neuronal and non-neuronal nuclei. We chose these thresholds as the minimal level that produced clear cluster separation (Figure 1E). A total of 31,258 genes were identified, with an average of 3,200 per nucleus. We performed standard log-normalization and a variance stabilizing transformation prior to finding anchors, and identified variable features individually for each monkeys’ data set using Seurat’s *FindVariableFeatures* function with number of features set to 7000. Next, we identified anchors using the *FindIntegrationAnchors* function with default parameters and passed these anchors to the *IntegrateData* function. This returned a Seurat object with an integrated expression matrix for all nuclei. We scaled the integrated data with the *ScaleData* function, ran PCA using the *RunPCA* function, and visualized the results with UMAP. We clustered the cells based on Louvain clustering and chose resolution that reflected the major cell classes of striatum including D1- and D2-MSNs, interneurons and astrocytes. We annotated the clusters based on feature plots of well-known marker genes (Gokce et al., 2016; Saunders et al., 2018; Zhang et al., 2014) and verified the identities of cell clusters by using the hypergeometric test to compare differentially expressed genes for each cluster to markers from single cell rodent studies (Grillner and Robertson, 2016). Given the robust conservation of major cell types, the rodent markers were sufficient for annotation (Grillner and Robertson, 2016). We converted rodent genes to rhesus macaque genes by BioMart Ensemble, keeping one-to-one orthologs only (Khrameeva et al., 2020; Yin et al., 2020).

In order to analyze MSNs in detail, we further analyzed a subset of the dataset that consisted of only MSN nuclei which were isolated according to the expression of several well-known marker genes including *PP1R1B*, *BCL11B*, and *PDE1B*. In order to balance the number of nuclei per sample, we randomly sampled MSN and MSN-like nuclei from monkey P to levels of monkey F (~2,800 nuclei for monkey F, 1 replicate; and ~5,600 nuclei for monkey P, 2 replicates). We then re-calculated principal components (PCs) and performed UMAP dimensionality reduction on the first 15 PCs. We used Louvain clustering to cluster the MSNs and MSN-like nuclei. We chose a resolution that reflected the main functional divisions in the striatum, including direct and indirect pathways, striosome and matrix compartments, and core and shell territories. To explore the functional roles of these cell types, gene enrichment analysis was run using gprofiler (Raudvere et al., 2019) with the following GO databases: Biological Process, Molecular Pathway, KEGG, and Human Phenotype ontology. We did hierarchical clustering using the *BuildClusterTree* function. To verify the validity of hierarchical clustering, we calculated the cosine similarity for nuclei within and between clusters based on PCA space. We used a permutation test on the cosine similarity between pairs of clusters. We randomly shuffled the nuclei to mask the nuclei identity and recalculated cosine similarity. We repeated these more than 10,000 times and used within group and between group variance ratio to determine a *P* value. To compare the cell types between monkey and mouse MSNs, we first generated an orthologous gene list between mouse and rhesus macaque from BioMart Ensemble with one-to-one orthologous genes (Khrameeva et al., 2020; Yin et al., 2020) used for downstream analysis. We integrated our MSNs and MSN-like nuclei from the two monkeys with MSNs types in Stanley et. al. (Stanley et al., 2020) based on the homologous genes using the similar above Seurat integration method.

### FISH image quantification

We quantified expression using high resolution images collected using a Nikon Eclipse T*i*2 (40x objective with or without additional 2x built in objective) or an Olympus IX83 under 40x or 63x magnification. For comparison between different regions, we scanned images using the same settings. To quantify cells expressing *DRD1* and *DRD2* in the caudate and putamen (Figure 3D), we chose ten representative areas from each striatal region (Figure 3C). We adjusted the threshold of the nuclei images and converted them to black and white using the “Make Binary” function in ImageJ. We then filled the holes using the “Fill Holes” function and separated overlapping nuclei with the “Watershed” function. We identified the number and regions for nuclei using the “Analyze Particles” function and added the nuclear contours to “ROI Manager”. We then opened the *DRD1* and *DRD2* images and identified the *DRD1* and *DRD2* grains using the “Find Maxima” function in ImageJ and then obtained binary images with a single pixel for each local maxima by choosing output type of “Single points”. We measured the integrated intensity of *DRD1* and *DRD2* above each nucleus using the “Measure” function within the “ROI Manager”. The number of grains in each nucleus is equal to the integrated intensity divided by 255. We considered cells containing three or more grains above the nucleus as positive for that gene. We chose this threshold because it provides clear separation from the background. We quantified the total number of *DRD1*-positive and *DRD2*-positive cells for each striatal region of interest (ROI) in each section and counted positive cells on three rostro-caudal sections. We calculated cell density as the number of positive cells divided by total number of nuclei in the area. We used a similar method to quantify the cell density in *CPNE4* and *RXFP1* clusters (Figure S5E) except that we drew ROIs for the whole *CPNE4* and *RXFP1* clusters after separating overlapping nuclei with the “Watershed” function. As control, we drew ROIs of roughly similar size to the *RXFP1* or *CPNE4* clusters in nearby regions and quantified the cells expressing *DRD1*.

To quantify *DRD1* and *DRD2* grains in D1/D2-hybrid cells, D1- and D2-MSNs (Figure 3H), we scanned high resolution images for *RXFP1*-positive cells in triple stained *DRD1*, *DRD2*, and *RXFP1* sections. The majority of *RXFP1*-positive cells expressed both *DRD1* and *DRD2* in dorsal striatum. We quantified the number of grains for *DRD1* and *DRD2* in these D1/D2-hybrid cells as well as adjacent normal D1 and D2 MSNs in ImageJ using similar methods as above except that we quantified the total grains in the cells instead of nuclei. To quantify *DRD1* and *DRD2* grains in D1/D2-hybrid cells, we first draw ROIs for *RXFP1* expressing cells and added the ROIs to the “ROI Manager” based on the *RXFP1* signal. We then opened the *DRD1* and *DRD2* images and identified the *DRD1* and *DRD2* grains using the “Find Maxima” function in ImageJ and then output binary images with a single pixel for each local maxima by choosing an output type of “Single points”. We measured the integrated intensity and calculated the number of grains for each ROI using the “Measure” function within the “ROI Manager”. To quantify *DRD1* and *DRD2* grains in D1- and D2-MSNs, we used the same method except that we drew ROIs for D1- and D2-MSNs. We confirmed the identity of D1- or D2-MSN instead of a D1/D2-hybrid cell by quantifying the *DRD2* or *DRD1* and *RXFP1* grain number less than three in D1- or D2-MSNs. We used a similar method to quantify *ARHGAP6* and *GREB1L* grains in the core and shell except that we first adjusted the threshold of the images to overexpose the *ARHGAP6* and *GREB1L* signals to guide the ROI drawing. To quantify *OPRM1* grain numbers in *RXFP1* and *CPNE4* islands and nearby D1-MSNs, we labeled *DRD1* and *OPRM1* on one section and *DRD1*, *RXFP1* and *CPNE4* on an adjacent section because of overlapping channels in the *OPRM1* and *RXFP1* probes. The locations of *RXFP1* and *CPNE4* islands in the *OPRM1* and *DRD1* double stained sections were determined by adjacent section labeled with *RXFP1* and *CPNE4*. The quantification of *OPRM1* grains numbers was similar to *ARHGAP6* or *GREB1L* quantification except that we used overexposed *DRD1* to draw ROIs for each cell.

To quantify nuclei size for *RXFP1* and *CPNE4* islands (Figure S5F), we first scanned high resolution images for *DRD1*, *RXFP1*, and *CPNE4* triple labeled sections. We performed automatic quantification of the areas of nuclei using ImageJ: 1) adjust the threshold and convert images to black and white using the “Make Binary” function; 2) fill the holes using the “Fill Holes” function; 3) separate overlapping nuclei with the “Watershed” function; 4) then draw the ROI and automatically detect the areas of nuclei using the “Analyze Particles” function. To produce a more accurate quantification of the nuclei size, we chose to randomly sample dozens of cells expressing *DRD1* in each island per section due to high packing density of the nuclei in ICjs (Figure S5E). We calculated the area of nuclei by the similar method except that we drew an area of interest around individual nucleus. Three sections in total were quantified and the area for each nucleus was normalized to the control D1-MSNs in each section.

### D1 islands mapping

In order to map the distribution of D1-RXFP1 and D1-CPNE4 islands, we triple labeled six sections spaced at 750 μm intervals with probes against *DRD1*, *RXFP1*, and *CPNE4* in monkey B. We scanned the whole sections and ran a custom CellProfiler pipeline (Carpenter et al., 2006) to determine the spatial distribution of nuclei and signal intensity of individual channel for each nucleus. We marked regions as D1-RXFP1 (D1-CPNE4) that had a majority of cells double-labeled with *DRD1* and *RXFP1* (*CPNE4*). To confirm that these islands were D1 exclusive, we double-labeled sections with *DRD1* and *DRD2* adjacent to the above six sections. We confirmed that these islands are exclusively the D1 clusters from the similar CellProfiler analysis. We did the similar process for eight sections spaced at 600 μm intervals for monkey K.

## DATA AND SOFTWARE AVAILABILITY

We deposited our data to GEO accession GSE167920. Go to https://www.ncbi.nlm.nih.gov/geo/query/acc.cgi?acc=GSE167920 and enter token otkbakeofdapfih into the box. Code is available at https://github.com/pfenninglab/nhp_snrna_striatum_analysis and https://github.com/rewardlab/NHPstriatum.

## SUPPLEMENTAL INFORMATION

Supplemental Information includes seven figures and three tables.

## Supplemental Figures

**Figure S1.**
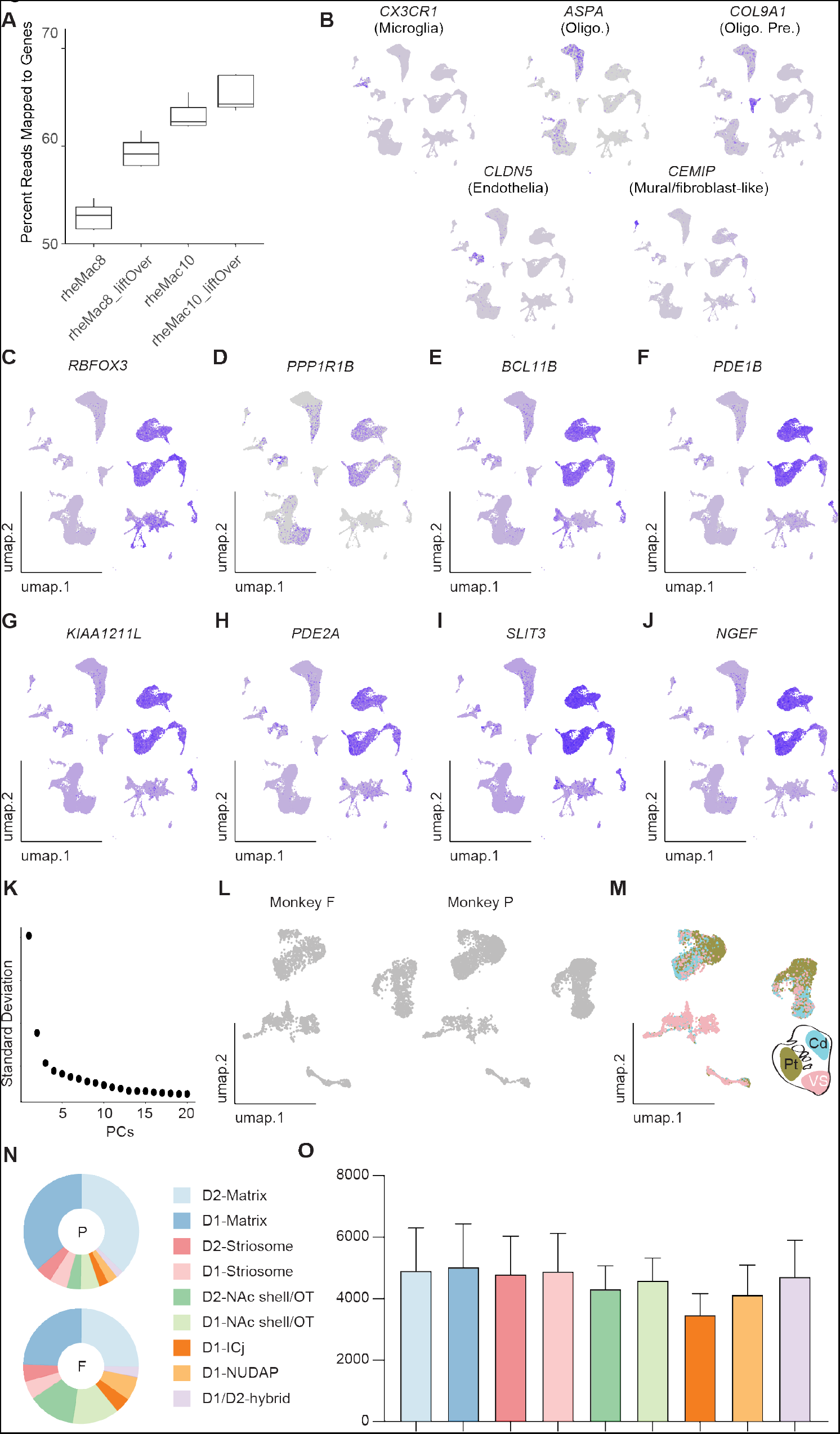
Read Mapping, Feature Plots, and Gene Expression Counting, Related to Figure 1 and 2. (A)Comparison of gene mapping rate using four different gtf in our alignment of reads to macaque genome. We used UCSC liftOver tool to liftOver the human transcriptome gtf file onto the rheMac8 or rheMac10 genome to get liftOver version rheMac8_liftOver and rheMac10_liftOver, respectively. (B)Feature plots of marker gene expression for major cell classes in striatal nuclei. (C)Feature plot showing the expression of *RBFOX3*, which labels all neurons. (D-J) Feature plots of gene expression for well-known MSN markers, *PPP1R1B* and *BCL11B*, *PDE1B* and new MSN-specific markers, *KIAA1211L*, *PDE2A*, *SLIT3*, *NGEF*. (K)Standard deviation in different principal components (PCs). The first 15 PCs account for majority of the variation. (L)Side by side monkey comparison for MSNs in UMAP coordinates. (M)UMAP visualization of MSNs colored by three regions. (N)Distribution of relative proportion of each MSN type in monkey P and F. (O)Gene expression across nine MSN types. Error bars indicate standard deviation across two monkeys.

**Figure S2.**
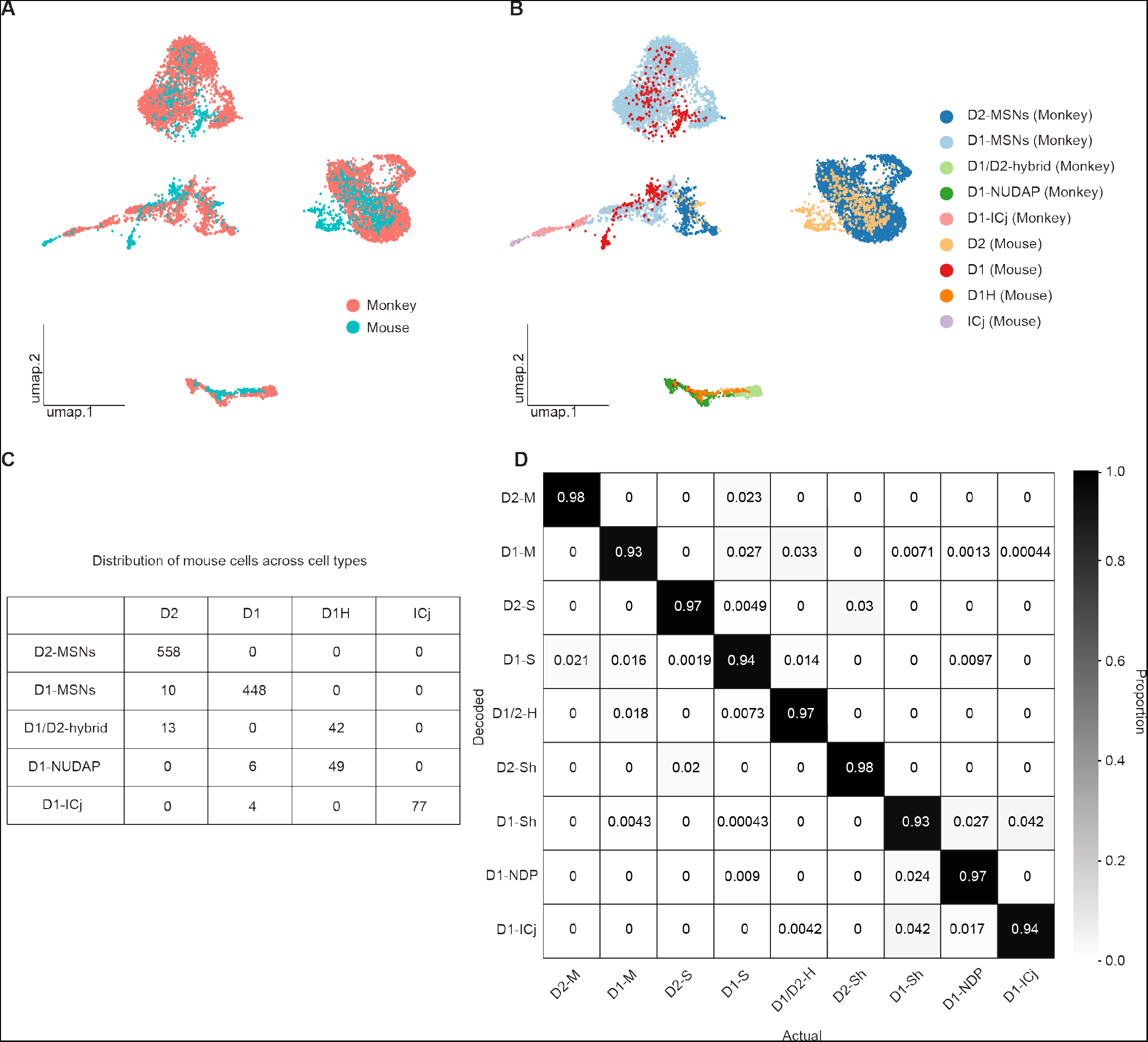
Monkey and Mouse MSN Comparison and Confusion Matrix Among MSN types, Related to Figure 2. (A)Integration of monkey and mouse MSNs in UMAP coordinates. (B)UMAP visualization of monkey and mouse MSN cell types. To make a comparable comparison, Monkey D1 striosome, matrix and shell/OT were combined as D1-MSNs (monkey). Similarly, monkey D2 striosome, matrix and shell were combined as D2-MSNs (monkey). (C)Combined monkey and mouse MSNs were re-annotated to (D2-MSNs, D1-MSNs, D1/D2-hybrid, D1-NUDAP and D1-ICj) and the number of cells per mouse MSN cell type that fell within the re-annotated clusters was quantified. (D)The accuracy rate (numbers within the grid) between SCCAF decoded cell type and actual cell type.

**Figure S3.**
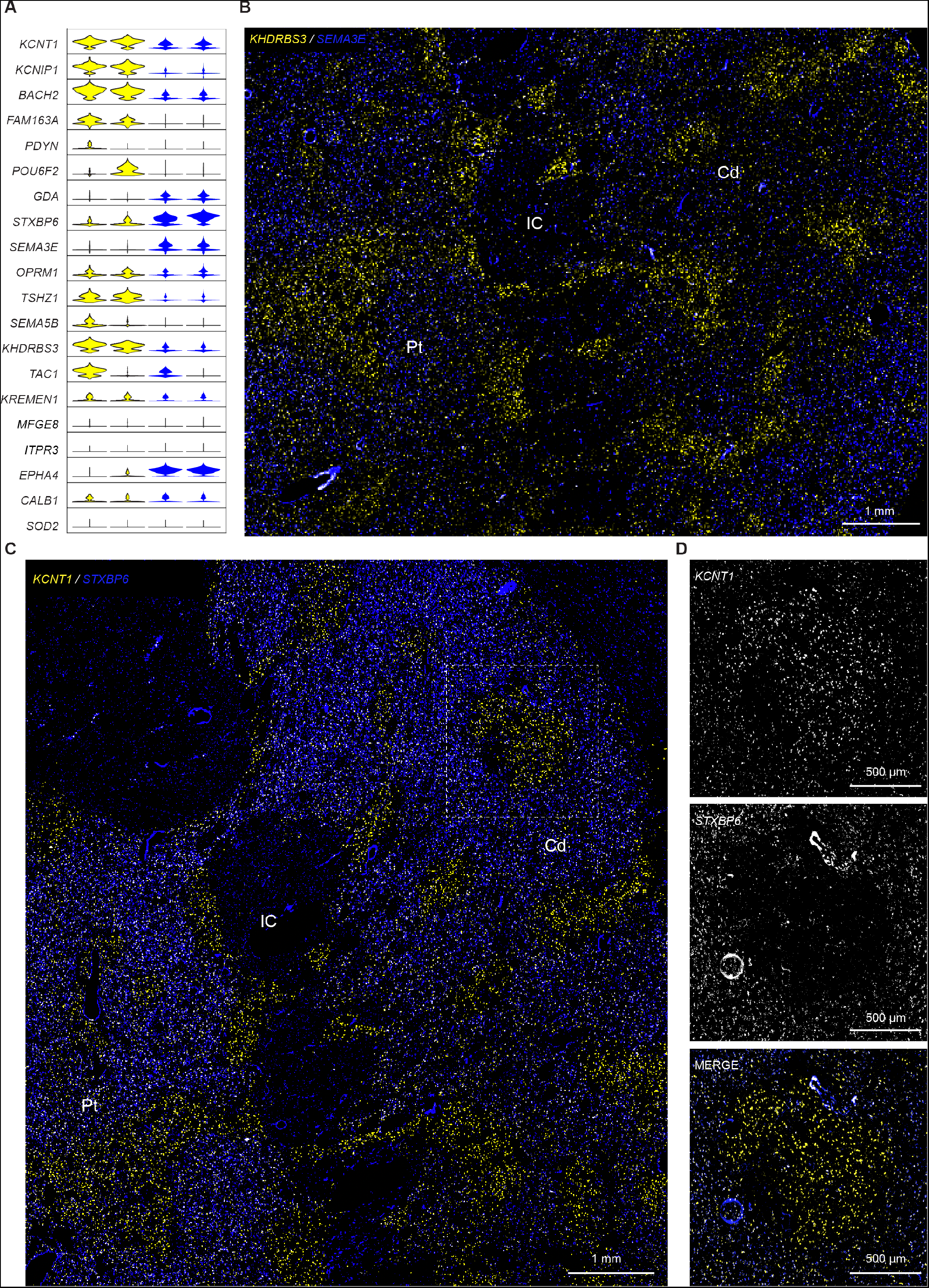
Striosome and Matrix Labeled by FISH Markers, Related to Figure 4. (A)Violin plot showing our identified and published striosome and matrix markers. From left to right columns are D1 striosome, D2 striosome, D1 matrix and D2 matrix. *PDYN* is a specific D1 striosome marker and *POU6F2* is a specific D2 striosome marker. (B)FISH labeling of *KHDRBS3* (yellow) and *SEMA3E* (blue) showing clear striosome and matrix distinction. (C)FISH labeling of *KCNT1* (yellow) and *STXBP6* (blue) showing striosome and matrix compartmentation. Cd: caudate, IC: internal capsule, Pt: putamen. (D)Detail of the white square in C. Scale bars are indicated on the images.

**Figure S4.**
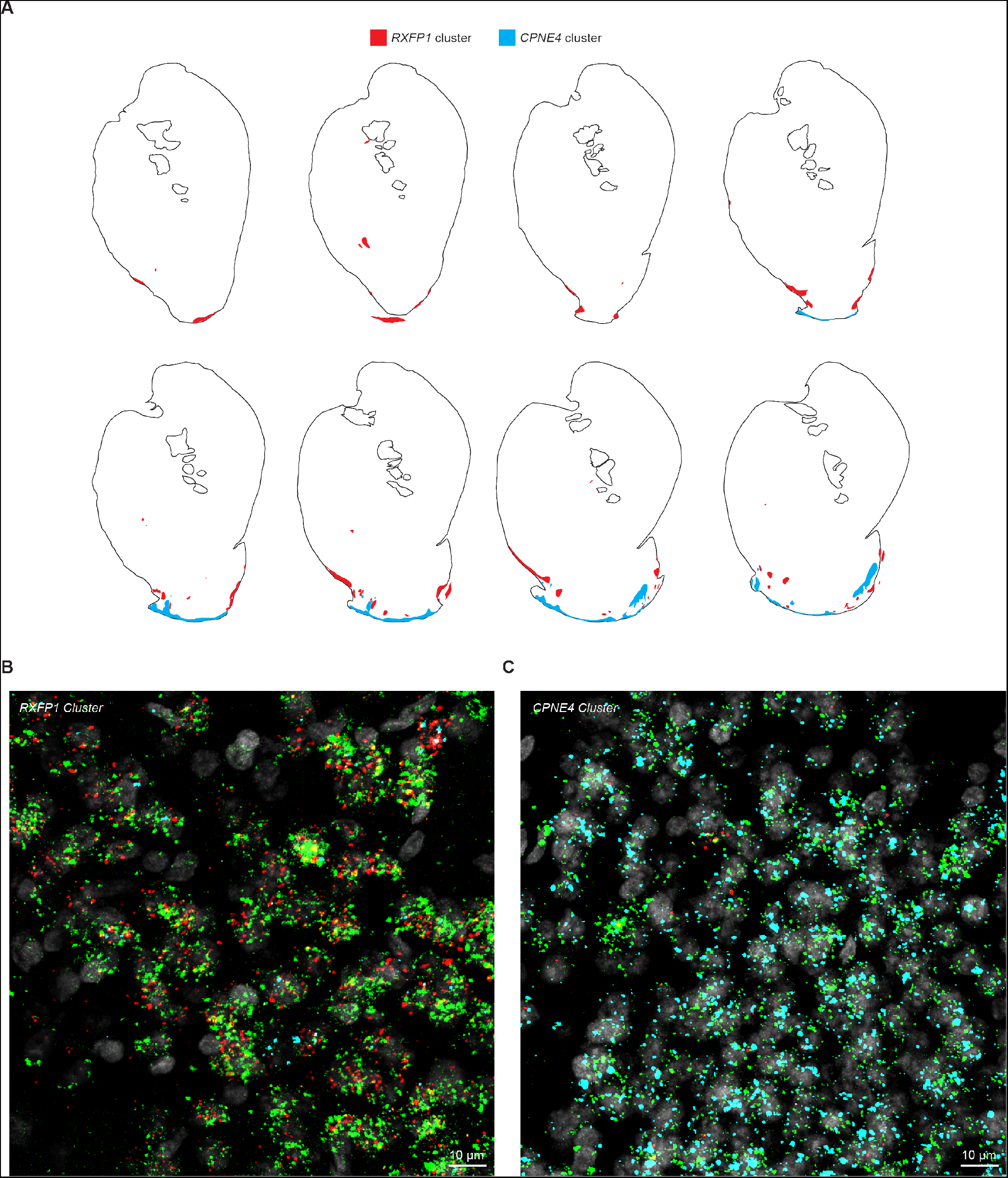
*RXFP1* and *CPNE4* Cluster Distribution in a Second Monkey, Related to Figure 6. (A)Distribution of *RXFP1* and *CPNE4* clusters across eight rostral-caudal regions identified by multichannel FISH in a second monkey K. (B)FISH stain of *DRD1* (green), *RXFP1* (red) and *CPNE4* (cyan). Nuclei were labeled by DAPI (grey). This image shows a representative *RXFP1* cluster in the ventral striatum. (C)FISH stain of *DRD1* (green), *RXFP1* (red) and *CPNE4* (cyan). Nuclei were labeled by DAPI (grey). This image shows a representative *CPNE4* cluster in the ventral striatum.

**Figure S5.**
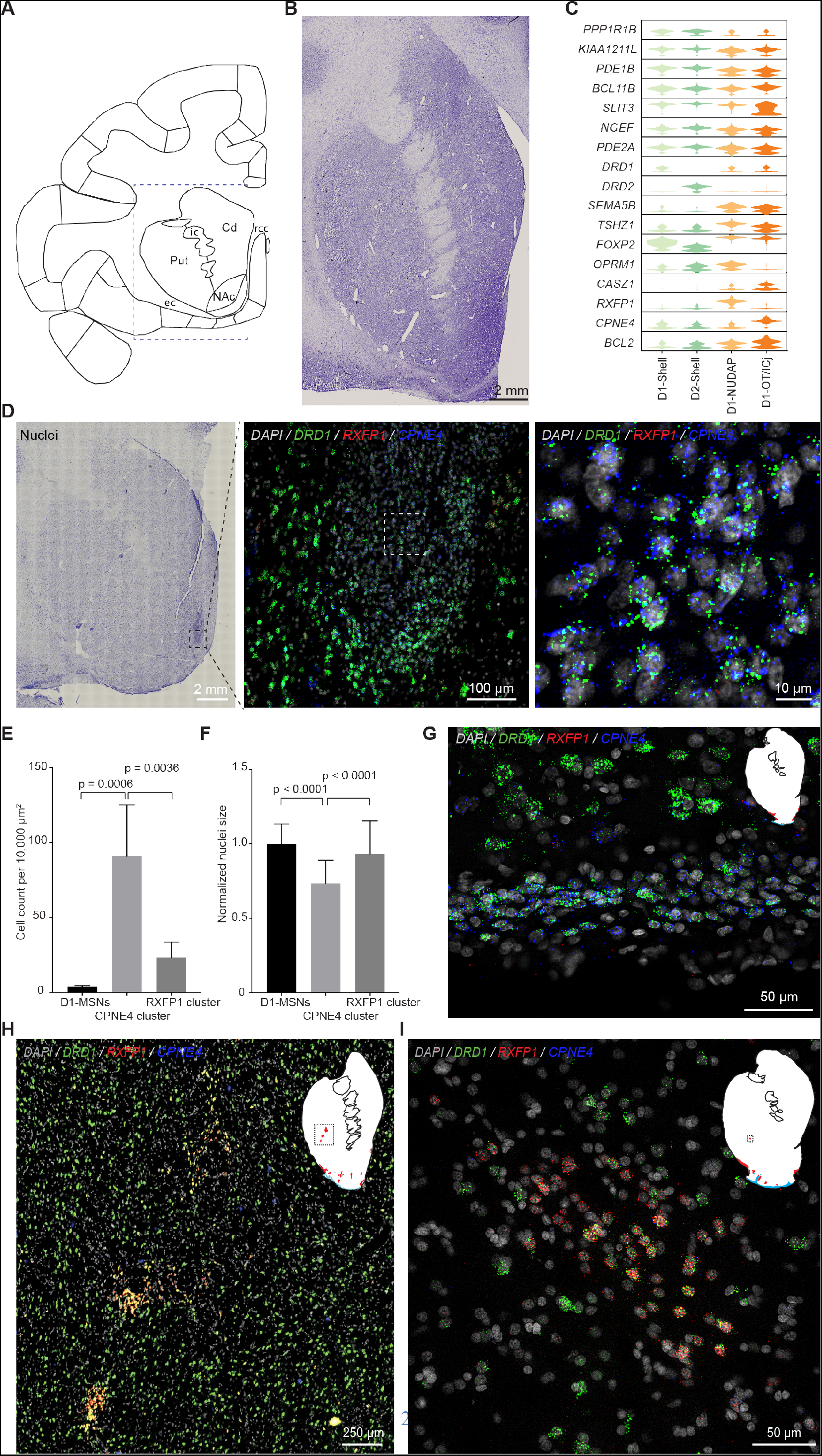
Comparing Nissl Section with Paxinos Atlas and ***RXFP1*** and ***CPNE4*** Cluster Mapping, Related to Figure 6. (A)Paxinos atlas shows the ventral extent of the external and extreme capsules loop around the ventral portion of the NAc and connects to the rostrum of the corpus callosum. Cd: caudate, Put: putamen, ec: external capsule, NAc: nucleus accumbens, rcc: rostrum of the corpus callosum, ic: internal capsule. (B)Nissl stain of one section. This Nissl image corresponds to the blue dashed box on the left. (C)Violin plots of relevant marker genes in ventral striatum MSN types. (D)Nissl stain of a section (left) and triple labeling of *DRD1* (green), *RXFP1* (red) and *CPNE4* (blue) in an adjacent section (middle and right) shows that *CPNE4* labels major island of Calleja. Nuclei labeled by DAPI (grey). Right image is the enlarged image from the boxed region in the middle image. (E)Quantification of cell density of neurons expressing *DRD1* in *RXFP1* and *CPNE4* clusters and nearby MSNs. One-way ANOVA with Bonferroni post hoc test was used for statistical analysis. Error bars are SD across 4 sections. (F)Quantification of nuclei size of neurons expressing *DRD1* in *RXFP1* and *CPNE4* clusters and nearby D1-MSNs. Nuclei size was normalized to the mean area size of regular D1-MSNs in each section. One-way ANOVA with Bonferroni post hoc test was used for statistical analysis. Error bars are SD across 49 cells from three sections. (G)FISH labeling of *DRD1* (green), *RXFP1* (red) and *CPNE4* (blue) in a section. Nuclei labeled by DAPI (grey). Inset: white area indicates striatum and the dashed black box highlights the area shown in the FISH image. The bottom *CPNE4* cluster mapped to AON. (H)FISH stain of *DRD1* (green), *RXFP1* (red), and *CPNE4* (blue) in one of section from monkey B. Inset: white area indicates striatum and the dashed black box highlights the area shown in the FISH image. (I)FISH stain of *DRD1* (green), *RXFP1* (red), and *CPNE4* (blue) in one of section from monkey K. Inset: white area indicates striatum and the dashed black box highlights the area shown in the FISH image.

**Figure S6.**
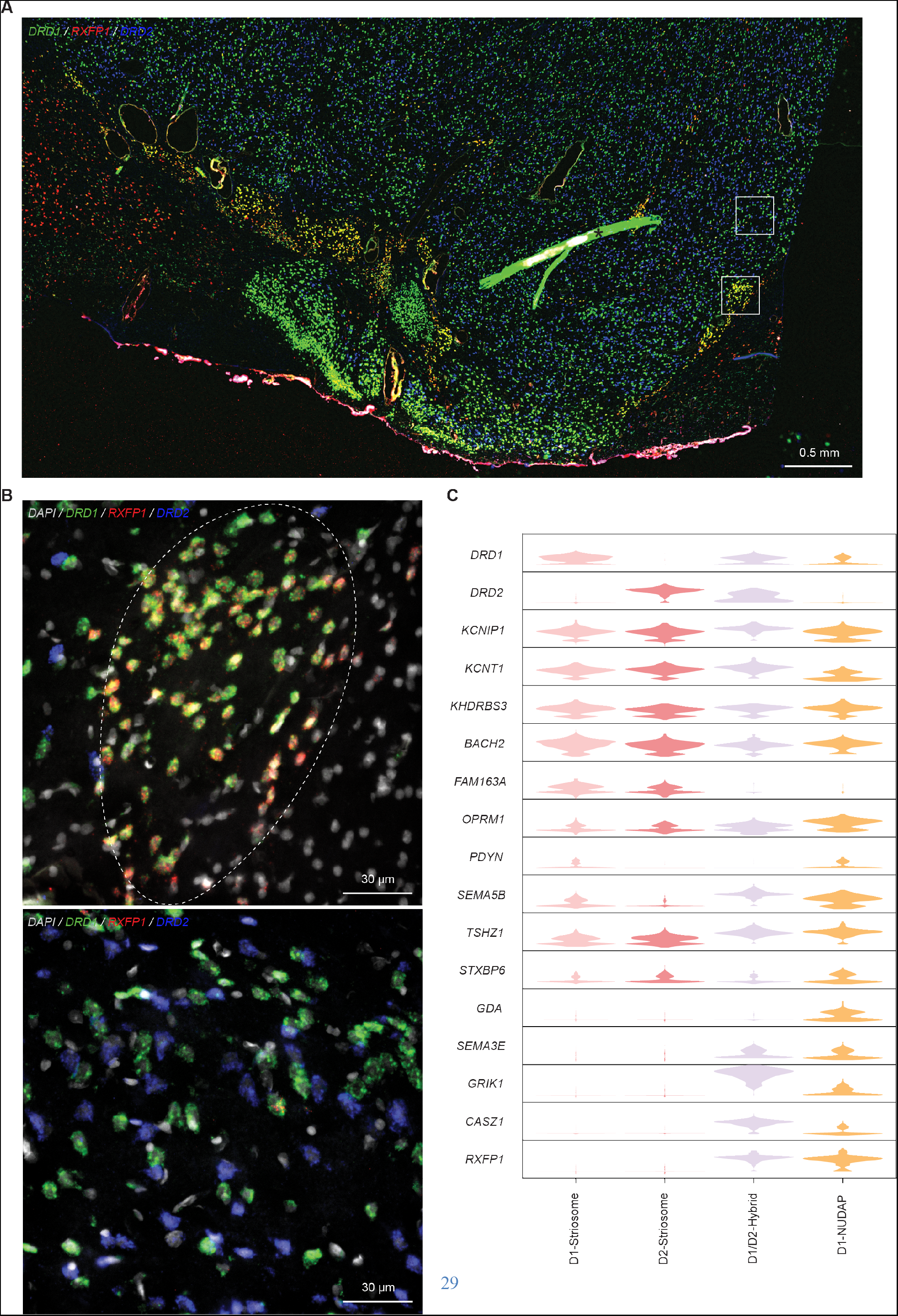
*RXFP1* Cluster and Violin Plots of Relevant Genes in Striosome, D1/D2-Hybrid and D1-NUDAP Cells, Related to Figure 7. (A)Triple FISH labeling of *DRD1* (green), *RXFP1* (red) and *DRD2* (blue) shows that *RXFP1* islands do not express *DRD2*. Black asterisk indicates staining artifact probably from dust. The artifact could be easily differentiated from real signals because there was no individual grain inside the artifact. (B)High resolution images from the boxed regions in A. (C)Violin plots of relevant marker genes in D1-striosome, D2-striosome, D1/D2-hybrid, and D1-NUDAP cells.

**Figure S7.**
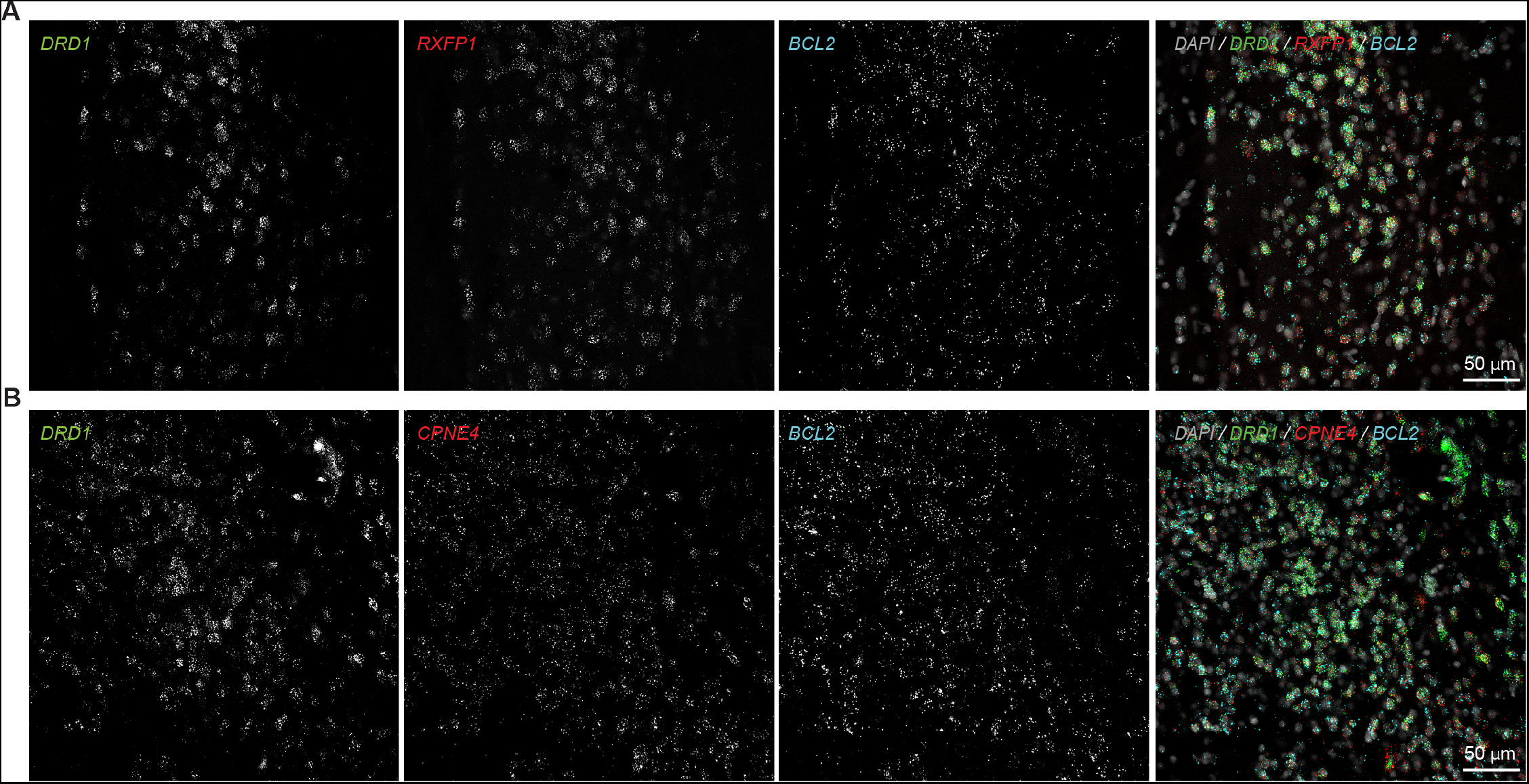
D1-NUDAP and D1-ICj Cells Are Both Labeled by *BCL2*, Related to Figure 7. (A)Triple FISH labeling of *DRD1* (green), *RXFP1* (red) and *BCL2* (cyan) shows that *RXFP1* islands express neuronal immature marker *BCL2*. (B)Triple FISH labeling of *DRD1* (green), *CPNE4* (red) and *BCL2* (cyan) shows that *CPNE4* islands express neuronal immature marker *BCL2*.

